# A SIRT5-induced metabolic switch underlies chemoresistance and ATR checkpoint dependence in triple-negative breast cancer

**DOI:** 10.64898/2026.04.07.716852

**Authors:** Zuen Ren, Tiziano Bernasocchi, Kiran Kurmi, Chenxu (Vincent) Guo, Kevin Jiang, Eric Zaniewski, Garrett Lam, Kazi N. Islam, Shakchhi Joshi, Xin Li, Ilze Smidt, Robert Morris, Bryce Ordway, Veerle I. Bossuyt, Gary X. Wang, Shinn-Huey S. Chou, Lee Zou, Ioannis Sanidas, Laura M. Spring, Michael Lawrence, Esther Rheinbay, Wilhelm Haas, Raul Mostoslavsky, Marcia C. Haigis, Leif W. Ellisen

## Abstract

Chemoresistance is the leading cause of poor prognosis in triple-negative breast cancer (TNBC), yet the underlying mechanisms remain unknown. To reveal metabolic drivers of de novo chemoresistance in TNBC, we analyzed pretreatment primary tumor biopsies, employing quantitative proteomics and metabolomics. Chemoresistant TNBCs exhibit hallmarks of oxidative phosphorylation (OXPHOS) and altered nucleotide metabolism linked to overexpression of the mitochondrial sirtuin, SIRT5. Through gain- and loss-of-function studies and stable isotope tracing, we demonstrate that SIRT5 induces a coordinated metabolic switch that redirects glycolysis to the pentose phosphate pathway, thereby augmenting nucleotide pools, while enhancing glutaminolysis to support OXPHOS. Mechanistically, SIRT5 enhances conversion of 6-phospho-D-gluconate to ribulose-5-phosphate through demalonylation of 6-phosphogluconate dehydrogenase (6-PGD), and coordinately activates oncogenic c-MYC to promote glutamine utilization and dependence. Concurrently, SIRT5-induced nucleotide deregulation induces replication stress and hypersensitivity to ATR checkpoint activation, and ATR inhibition synergistically reverses chemoresistance in TNBC. Thus, elevated SIRT5 orchestrates a coordinated metabolic switch to expand nucleotide pools and drive chemoresistance, while producing ATR checkpoint dependence that represents a metabolic vulnerability of SIRT5-overexpressing TNBC.

**Graphical Abstract:** 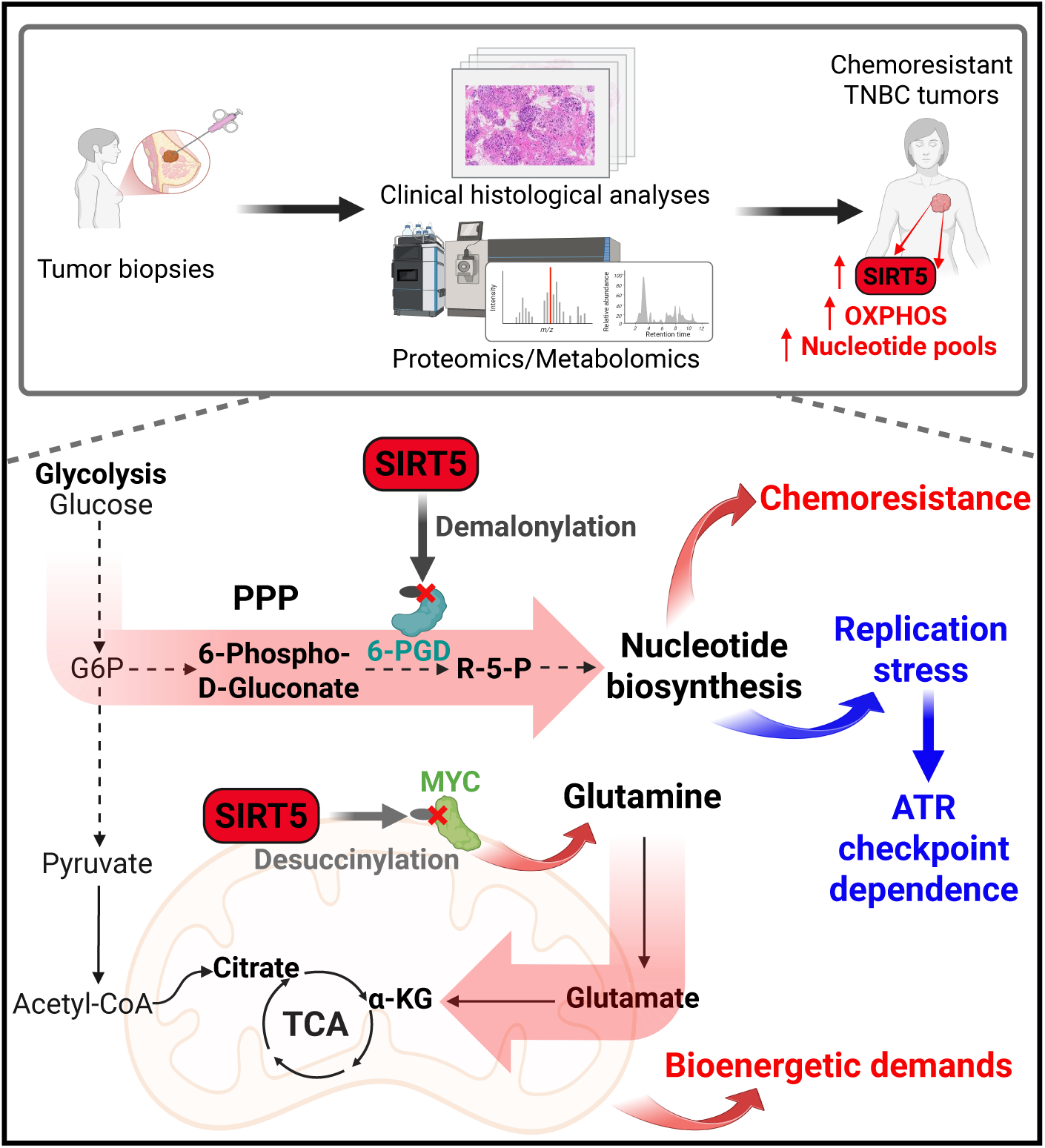

## Introduction

Triple-negative breast cancers (TNBCs)—characterized clinically by the absence of estrogen receptor (ER) and progesterone receptor (PR) expression and human epidermal growth factor receptor 2 (HER2) gene amplification—frequently manifest *de novo* drug resistance (1). Inter-tumoral heterogeneity and the lack of actionable driver mutations have hampered the advancement of targeted therapies for TNBC, leaving cytotoxic chemotherapy with or without immunotherapy as a primary treatment modality. While pre-operative chemotherapy that incorporates anthracycline and taxane-based regimens is a standard treatment approach for primary TNBC, it often fails to achieve complete tumor eradication, known as pathological complete response (pCR), owing in part to the inherent chemoresistant characteristics of these tumors. Since 70% patients whose tumors fail to undergo pCR experience a high rate of relapse, there is an urgent need to deepen our understanding of the intrinsic features of treatment-naive tumors that contribute to *de novo* chemoresistance in TNBC, to enable effective therapeutic strategies for TNBC (2).

Tumor metabolic reprogramming plays a crucial role in conferring inherent resistance to chemotherapy in cancer cells (3-5). In a context- and tumor-specific manner, alterations of crucial metabolic pathways that support rapid cell growth and survival not only drive tumorigenesis but also mediate treatment resistance and metastasis, collectively conferring adverse clinical outcomes (6-10). Understanding the detailed mechanisms and consequences of metabolic reprogramming that govern drug resistance can pave the way for new therapeutic approaches and better treatment outcomes. Diverse features of aberrant tumor metabolism include the role of altered glucose, glutamine, lipid, nucleotide, and amino acid metabolism in driving breast cancer progression and treatment response, yet the identification of specific mediators and biochemical mechanisms linking altered tumor metabolism to *de novo* treatment resistance has remained largely elusive (11-15).

Major obstacles to understanding physiologically relevant tumor metabolism in drug resistance include context specificity and generally poor accuracy of mechanisms identified primarily through cell-based studies (16, 17). Consequently, we sought to employ analysis of primary human tumors in a well-defined therapeutic setting as our initial discovery approach. Utilizing meticulously collected breast tumor biopsies from treatment-naive breast cancer patients, we conducted proteomic, metabolomic, and clinical histological analyses to investigate the intrinsic metabolic characteristics of diverse tumor subtypes. Our initial observations revealed, somewhat surprisingly, similarity between benign fibroadenomas and hormone receptor-positive (HR+) tumors, while TNBC tumors exhibited distinct metabolic profiles. We then carried out a detailed comparison of TNBC tumors that underwent pCR following pre-operative chemotherapy versus chemoresistant tumors that did not. We observed that mitochondrial sirtuin SIRT5 serves as the nexus of metabolic rewiring and is overexpressed in chemoresistant tumors. SIRT5 is an atypical sirtuin in that it lacks robust deacetylase activity and primarily functions to remove succinyl, malonyl, and glutaryl modifications from lysines on its substrates in mitochondria and throughout the cell, thereby regulating multiple important metabolic pathways (18-23). Here, we define the mechanisms by which high levels of SIRT5 in TNBC coordinately deregulate cellular metabolism to expand nucleotide pools that underlie both chemoresistance and replication stress checkpoint sensitivity, thus creating a therapeutically actionable metabolic vulnerability in these refractory tumors.

## Results

### Distinct metabolic and proteomic characteristics of primary breast tumors

We first sought to define the distinct metabolic profiles of benign versus malignant primary breast tumors and to compare breast cancer subtypes. We obtained additional snap-frozen core needle biopsies with informed consent from patients undergoing routine clinical breast biopsies for diagnostic purposes (24). The tissue was frozen within two minutes of collection to preserve metabolite and proteome profiles. Based on standard clinicopathologic assessment, we selected 12 TNBC samples (grade 3), 12 HR+ invasive ductal carcinoma (IDC) samples (grade 3), and 12 benign fibroadenoma samples for comprehensive metabolomic, proteomic, and histological analyses (**Fig. 1A** and **Table S1-2**).

**Fig. 1.**
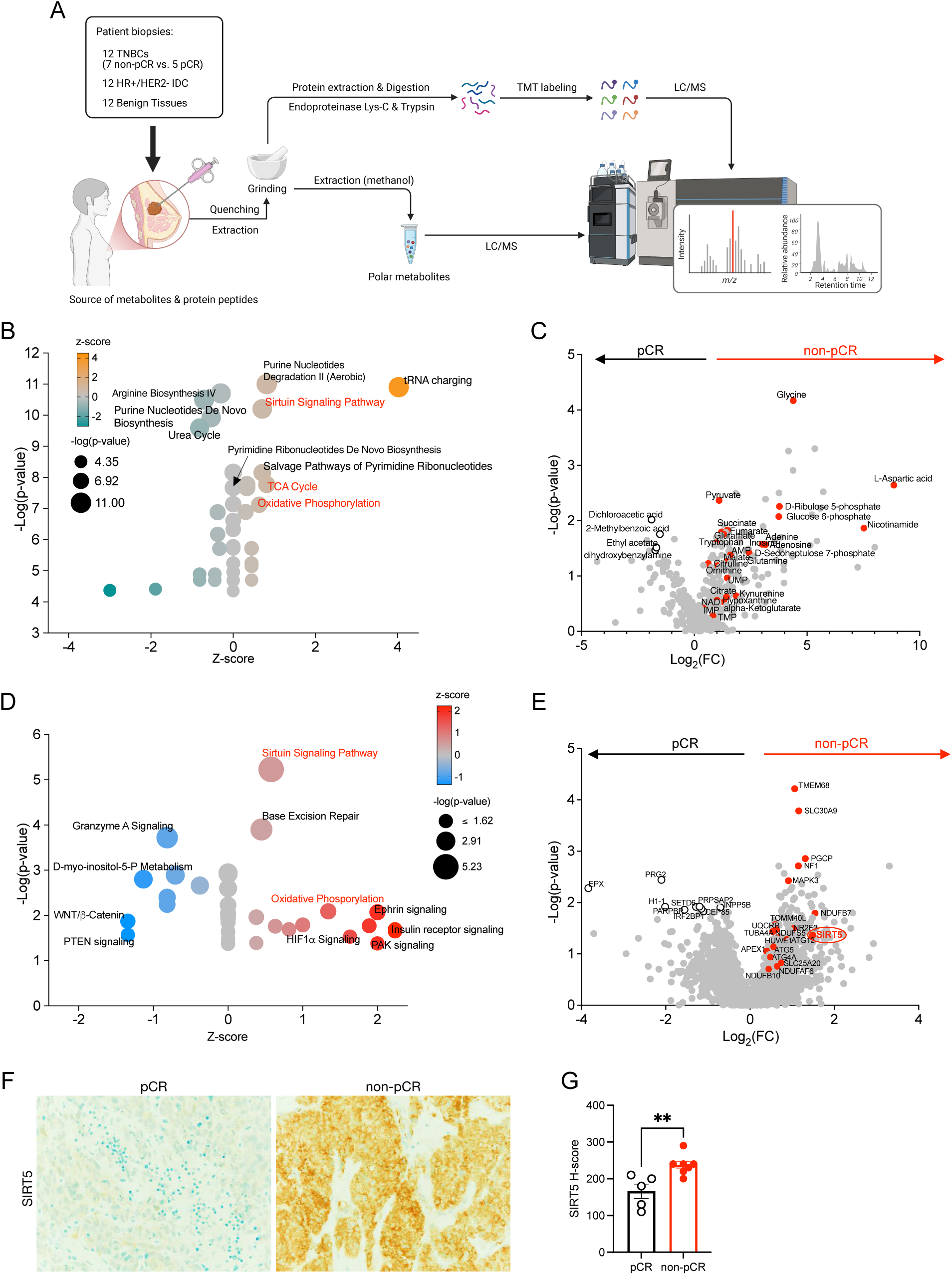
Proteomic, metabolomic, and clinical histological analyses uncover metabolic traits associated with de novo chemoresistance in TNBC. **A.** Schematic representation of the workflow displays quantitative high-throughput Liquid Chromatography-Mass Spectrometry (LC/MS) analysis of protein and metabolite profiles from TNBC (n = 12), HR+/HER2-invasive ductal carcinoma (IDC) (n = 12), and benign breast tissues (n = 12). **B.** Ingenuity Pathway Analysis (IPA) highlights significantly overrepresented canonical pathways in non-pathological complete response (non-pCR) TNBC tumors derived from metabolomics. Highly enriched pathways, including the Sirtuin signaling pathway, the tricarboxylic acid (TCA) cycle, and oxidative phosphorylation (OXPHOS), are highlighted in red. **C.** Volcano plot illustrates differential expression of metabolites between non-pCR versus pCR TNBC tumors; metabolites highlighted in red correspond to those associated with pathway dependencies indicated in panel B. **D.** IPA reveals overrepresented canonical pathways in non-pCR TNBC tumors based on proteomic data, including the Sirtuin signaling pathway and OXPHOS (in red). **E.** Volcano plot depicting the differentially expressed proteins between non-pCR and pCR TNBC tumors; proteins marked in red denote those associated with the Sirtuin signaling pathway and OXPHOS in panel D. **F-G**. Immunohistochemical (IHC) staining (F) for SIRT5 on matched formalin-fixed, paraffin-embedded (FFPE) samples of TNBC reveals significantly higher expression of SIRT5 in non-pCR tumors compared to pCR tumors, as quantified by the intensity of IHC staining (G). ** p-value < 0.01.

Each frozen core biopsy was cryopulverized and divided to enable both metabolite and proteome profiling (**Fig. 1A**). Principal component analysis (PCA) of targeted metabolites (n=155, see Methods and **Table S3**) revealed substantial overlap between fibroadenomas and HR+ breast cancers, while TNBC primary tumors exhibited a distinct metabolic profile (**Fig. S1A**). Hierarchical clustering of metabolite levels further illustrated the distinct profiles of TNBCs compared to HR+ IDC tumors and benign tumors (**Fig. S1B**). Notably, metabolites involved in glycolysis and the TCA cycle were highly enriched in both HR+ and TNBC primary tumors, particularly in TNBC tumors, and metabolites involved in O-glycosylation for cell-cell adhesion, such as N-acetylgalactosamine (GalNAc), were uniquely decreased in TNBC primary tumors as compared to both HR+ and benign breast tumors (**Fig. S1B**). The altered GalNAc expression has been used as a biomarker for early cancer detection and prognosis (25, 26). Furthermore, Ingenuity Pathway Analysis (IPA) of the differentially expressed metabolites between TNBC and benign tumors supported the notion that glycolysis and the TCA cycle are highly enriched metabolic pathways in TNBC primary tumors (**Fig. S1C**).

By contrast, proteomic profiling (n=7309 proteins detected, see Methods and **Table S4**) via Reversed-Phase Liquid Chromatography (RPLC) coupled with mass spectrometry (MS) did not reveal as pronounced a distinction between tumor types on PCA as the metabolite analysis (**Fig. S1D**). The most notable distinction upon hierarchical clustering of proteomes among the three tumor types was the relative homogeneity within the fibroadenomas and HR+ IDC tumor subsets (albeit with different sets of highly expressed proteins) compared to the marked inter-tumoral heterogeneity among TNBC tumors (**Fig. S1E**). Interestingly, we found that numerous proteins involved in extracellular matrix formation, such as LAMA, LAMB, LAMC, COL18A1, and COL15A1, were down-regulated in malignant primary breast tumors and more consistently in HR+ breast tumors, underscoring the importance of extracellular matrix proteins in breast cancer progression and treatment (**Fig. S1E**). In agreement with the metabolic pathways identified through metabolite profiling, IPA of the differentially expressed proteins between TNBC and benign tumors recapitulated a significant number of highly enriched metabolic pathways in TNBC primary tumors, including arginine biosynthesis, the superpathway of methionine degradation, glycolysis, and the TCA cycle (**Fig. S1F**). These findings suggest that metabolic signatures enriched in TNBC tumors are consistently recapitulated across multiple analytes. The distinct enrichment of glycolysis and the TCA cycle in TNBC tumors may allude to proposed models whereby a switch between aerobic glycolysis (Warburg effect) and OXPHOS allows environmental adaptations that support tumor progression and metastasis (27, 28).

#### Intrinsic metabolic properties of chemoresistant TNBC revealed by proteomics and metabolomics

To identify metabolic features associated with treatment response and long-term patient outcomes, we next focused on a comparison of pretreatment TNBC samples from patients whose tumors underwent pCR following subsequent preoperative (neoadjuvant) therapy versus those that did not (non-pCR) (see **Table S1**). All patients received systemic neoadjuvant chemotherapy, which primarily consisted of dose-dense adriamycin and cyclophosphamide, followed by paclitaxel (ddAC/T), before undergoing mastectomy or lumpectomy (see **Table S1**). We separately performed a detailed targeted and untargeted metabolomics analysis of these tumors, resulting in the profiling of 360 metabolites (**Table S5**). Additionally, a reanalysis of the proteomics data from TNBC primary tumors yielded a total of 6,278 proteins after data cleaning and filtering (see **Table S6**). IPA comparing the differentially expressed metabolites between non-pCR and pCR TNBC tumors indicated significant enrichment in pathways related to the TCA cycle, OXPHOS, sirtuin signaling, and nucleotide biosynthesis in non-pCR TNBC tumors (**Fig. 1B**). Enrichment was hallmarked by metabolites involved in these pathways, such as glycine, aspartic acid, pyruvate, succinate, fumarate, ribulose-5-phosphate, and glucose-6-phosphate (**Fig. 1C**). Correspondingly, IPA comparing the differentially expressed proteins between non-pCR and pCR TNBC tumors revealed significant enrichment of OXPHOS, sirtuin signaling, and base excision repair pathways at the protein level in non-pCR TNBC tumors (**Fig. 1D**). Importantly, both protein and metabolite analyses identified OXPHOS and Sirtuin signaling pathways as the top significantly enriched pathways in non-pCR tumors (see **Fig. 1B and 1D**). This finding is consistent with previous studies suggesting that a high OXPHOS-dependent energetic state contributes in some way to chemoresistance in TNBC (29-32).

Differential analysis of the proteomics data between non-pCR and pCR TNBC tumors identified SIRT5 as the only sirtuin family member that was significantly differentially expressed in non-pCR vs. pCR tumors (**Fig. 1E**). SIRT5 is a sirtuin family member localized primarily to the mitochondrial matrix. Unlike typical mitochondrial sirtuins SIRT3 and SIRT4, which function primarily as nicotinamide adenine dinucleotide (NAD)-dependent mitochondrial protein deacetylases, SIRT5 primarily removes succinyl and malonyl modifications from lysines on target proteins in both mitochondria and the cytosol (18-23). Emerging literature suggests that SIRT5 plays a role in regulating mitochondrial metabolism and OXPHOS, which may contribute in various contexts to drug resistance and tumor progression (33-37). To confirm the presence of SIRT5 in the profiled TNBC tumors, we assessed immunohistochemical (IHC) staining in the corresponding clinical formalin-fixed paraffin-embedded (FFPE) tumor specimens. We found that SIRT5 protein was statistically significantly higher in non-pCR tumors compared to pCR tumors, with high-level expression localized nearly exclusively in tumor cells (**Fig. 1F-G**). Collectively, these findings suggest the hypothesis that the elevated expression of SIRT5 in non-pCR TNBC tumors may underlie a high OXPHOS-dependent energetic state that could contribute to chemoresistance.

To investigate the chemoresistant phenotype in more detail, we next carried out an integrated analysis of metabolite and proteomic profiles of these primary TNBC tumors using the Metaboverse platform (38). Metaboverse is an interactive platform designed for the exploration of multi-omics data within the context of metabolic networks, utilizing information about reaction inputs, outputs, catalysts, and inhibitors (38). Consistent with the metabolic signatures identified in the IPA analysis, Metaboverse revealed similar top-enriched metabolic networks in non-pCR TNBC tumors, including the TCA cycle, Urea cycle, Nucleotide salvage, aspartate and asparagine metabolism, Purine catabolism, and Base excision repair (**Fig. S2A-F**). Overall, these findings demonstrate that the metabolic signatures enriched in non-pCR vs. pCR TNBC tumors can be consistently recapitulated across multiple analytes using distinct analytic approaches.

### Elevated SIRT5 expression and gene copy number gain/amplification are enriched in TNBC and associated with poor outcomes following chemotherapy

Analysis of SIRT5 mRNA expression and genomic alteration across breast cancer subtypes in The Cancer Genome Atlas (TCGA) and the Molecular Taxonomy of Breast Cancer International Consortium (METABRIC) databases (39, 40), indicated that TNBC tumors exhibit significantly higher levels of SIRT5 mRNA expression compared to non-TNBC tumors (**Fig. 2A and Fig. S3A**). We also assessed SIRT5 mRNA expression across different intrinsic breast cancer subtypes via PAM50 analysis. Consistently, SIRT5 mRNA expression was markedly elevated in the basal-like subtype (predominantly TNBC) relative to luminal A/B, HER2-enriched, and normal-like breast cancer subtypes (**Fig. 2B and Fig. S3B**). Unlike SIRT5, the other mitochondrial sirtuins SIRT3 and SIRT4 did not display significant differences across breast cancer subtypes (**Fig. S3C-F**). Together, these results indicate that SIRT5 is the only mitochondrial sirtuin significantly overexpressed in TNBC.

**Fig. 2.**
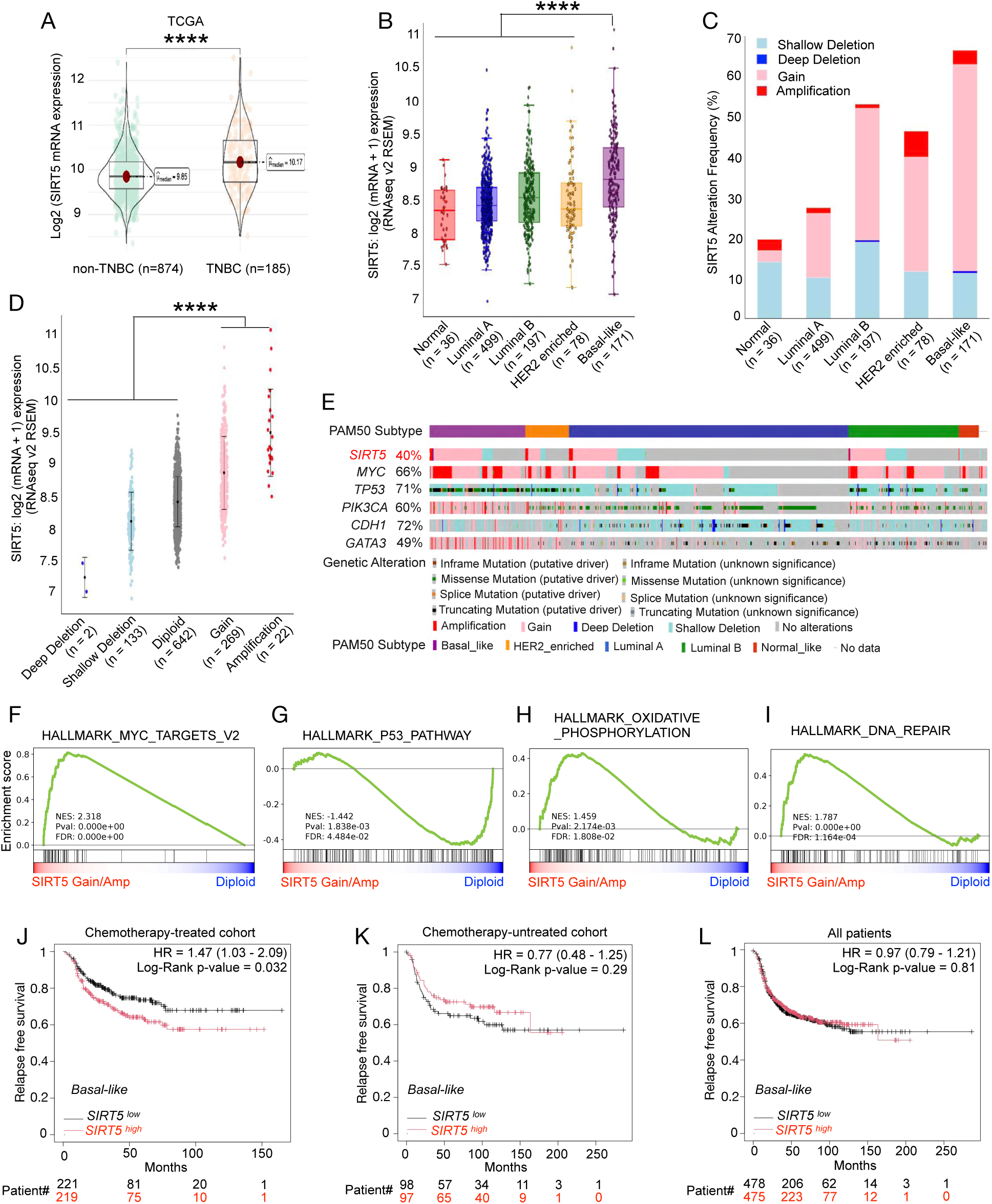
SIRT5 overexpression is associated with copy number gain and amplification and predicts chemoresistance in breast cancer. **A.** Violin plots show the distribution of SIRT5 mRNA expression levels between TNBC and non-TNBC samples from the TCGA database, highlighting a significant increase in TNBC. **B.** Box plots depict log2-transformed SIRT5 mRNA expression levels across breast cancer subtypes classified by the PAM50 gene expression signature, revealing the highest SIRT5 levels in basal-like tumors (primarily TNBC). **C.** Bar graphs illustrate the frequency of SIRT5 genetic alterations across different breast cancer subtypes, indicating the highest frequency of gains in basal-like tumors. **D.** Dot plots present the association between SIRT5 mRNA expression levels and the frequency of SIRT5 genetic alterations, establishing a connection between mRNA expression and genetic alterations across breast tumors. **E.** Oncoplot visualization presents the alteration frequency of commonly mutated genes across the different subtypes of breast cancer, revealing significant co-association of gains/amplification of SIRT5 and MYC (Fig. S3G). **F-I**. Gene Set Enrichment Analysis (GSEA) identifies hallmark pathways that are significantly enriched in breast tumors with SIRT5 gain/amplification (n = 291) compared to diploid tumors (n = 642), revealing enrichment of MYC, OXPHOS, and DNA repair signatures associated with SIRT5. (Data for panels A-I were sourced from the cBioPortal database: https://www.cbioportal.org/study/summary<u>?</u> <u>id=brca_tcga_pan_can_atlas_2018</u>.) **J-L**. Kaplan-Meier survival analysis demonstrates that high SIRT5 expression is significantly correlated with poorer relapse-free survival in chemotherapy-treated patients with basal-like breast cancer, whereas this correlation is absent in untreated breast cancer patients. (Data for panels J-L were derived from KM-plotter, which integrates gene expression and clinical data from GEO, EGA, and TCGA). **** p-value < 0.0001.

Next, we explored the correlation between SIRT5 mRNA expression and genomic alterations. SIRT5 showed frequent copy number gain and genomic amplification in breast cancer, particularly within the basal-like subtype (**Fig. 2C**). Approximately sixty percent of basal-like breast cancers exhibited SIRT5 gene amplification or copy number gain (**Fig. 2C**). Additionally, high SIRT5 mRNA levels were found to correlate with SIRT5 gene amplification and copy number gain in breast tumors (**Fig. 2D**). We further investigated the frequency of co-alterations in SIRT5 and other commonly altered genes across various breast cancer subtypes, including MYC, TP53, PIK3CA, CDH1, and GATA3 (**Fig. 2E**). We observed that SIRT5 gain/ amplification was significantly correlated with MYC amplification/copy gain and TP53 mutations (**Fig. S3G-H**), consistent with findings that high expression of MYC and mutations in TP53 are associated with worse outcomes in TNBC (41). Thus, SIRT5 overexpression in TNBC is driven by copy number gain and amplification and is associated with other genomic features of particularly aggressive TNBC.

These findings were further supported by the differential gene expression analysis we performed comparing breast tumors with SIRT5 gain/amplification to tumors diploid for SIRT5 (**Fig. S3I**). As anticipated, breast tumors harboring SIRT5 gain/amplification displayed significantly higher SIRT5 mRNA levels compared to SIRT5 diploid tumors (**Fig. S3I**).

Furthermore, Gene Set Enrichment Analysis (GSEA) revealed that MYC target genes were significantly enriched in tumors with SIRT5 gain/amplification compared to diploid tumors, while canonical TP53-induced pathways were negatively associated with SIRT5 gain/amplification (**Fig. 2F-G**). GSEA also identified significant enrichment in OXPHOS and DNA repair pathways in breast tumors with SIRT5 gain/amplification, consistent with our findings from metabolomic and proteomic comparison of non-pCR versus pCR TNBC tumors. Accordingly, Kaplan-Meier analysis demonstrated that elevated SIRT5 levels predict poorer relapse-free survival selectively in chemotherapy-treated basal-like breast cancers (**Fig. 2J**). Notably, this association was absent among untreated basal-like breast cancers (**Fig. 2K-L**) (42). This finding suggests that SIRT5 is linked specifically to chemoresistance rather than serving as a general prognostic indicator. Collectively, these results corroborate our data from the primary TNBC tumors, revealing SIRT5 as a hallmark of metabolic reprogramming and chemoresistance in these tumors.

### SIRT5 expression induces chemoresistance through its enzymatic activity

To probe the potential role of SIRT5 in TNBC chemoresistance, we examined SIRT5 expression across multiple TNBC cell lines and observed heterogeneous expression levels (**Fig. 3A**). Notably, the Hs578T TNBC cell line, which exhibits little or no SIRT5 expression, demonstrated higher sensitivity and an IC50 to doxorubicin and cisplatin nearly 10-fold lower compared to SIRT5-expressing TNBC lines (**Fig. 3A-B and Fig. S4A**). Moreover, a linear regression analysis showed a correlation between doxorubicin IC50 values and SIRT5 protein expression across these TNBC cell lines (**Fig. 3C**).

**Fig. 3.**
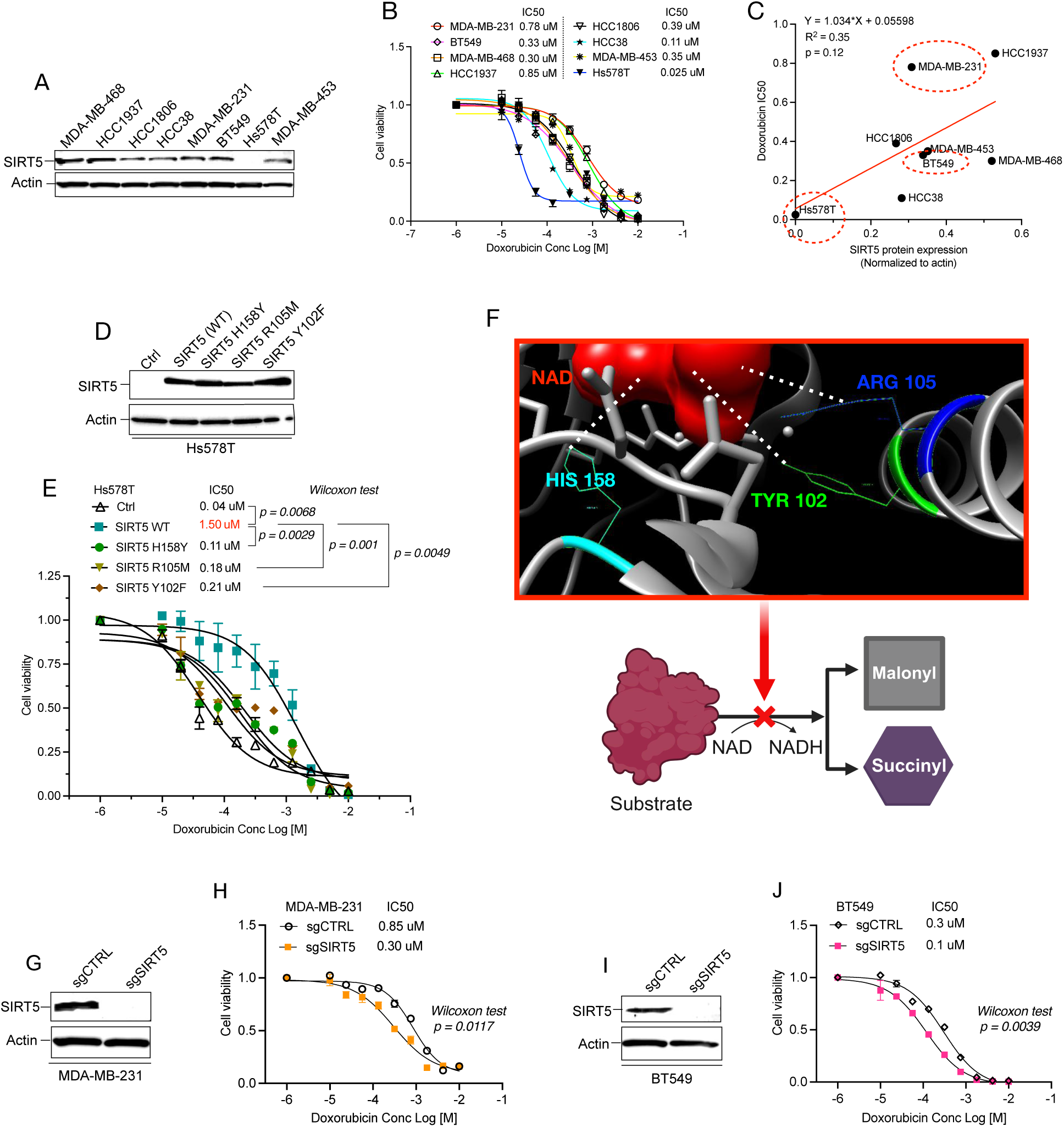
SIRT5 promotes chemoresistance in TNBC cells, dependent on its enzymatic activity. **A.** Western blotting shows SIRT5 and actin (control) expression levels across various TNBC cell lines. **B.** Dose-response curves show the sensitivity of TNBC cell lines to doxorubicin; corresponding IC50 values are indicated. **C.** A linear regression analysis presents the correlation of doxorubicin IC50 values with SIRT5 protein expression (normalized to actin) across various TNBC cell lines. **D.** Western blot analysis assesses the expression of SIRT5, SIRT5 mutants, and actin proteins in Hs578T. **E.** Dose-response curves demonstrate the effect of SIRT5 wild-type (WT) and mutant expression on doxorubicin sensitivity in Hs578T TNBC cells; IC50 values are indicated. **F.** The crystal structure of SIRT5 (PDB: 3RIY) displays the ternary structure of the ligand NAD interacting with key residues Tyr102, Arg105, and His158 in the catalytic domain. SIRT5 exhibits demalonylase and desuccinylase activities on its substrates. **G.** Western blots show SIRT5 and actin levels in MDA-MB-231 following introduction of sgSIRT5 or control. **H.** Dose-response curves show doxorubicin sensitivity of MDA-MB-231 cells with sgCTRL and sgSIRT5; IC50 values are indicated. **I.** Western blots show SIRT5 and actin levels in BT549 following introduction of sgSIRT5 or control. **J.** Dose-response curves show doxorubicin sensitivity of BT549 cells with sgCTRL and sgSIRT5; IC50 values are indicated.

To investigate directly the impact of SIRT5 expression on sensitivity to doxorubicin and cisplatin, we conducted both gain and loss-of-function studies. Expression of wild-type (WT) SIRT5 in Hs578T cells resulted in a marked increase in resistance to doxorubicin, with an IC50 value that was 37.5 times higher than that of control cells transfected with an empty vector (**Fig. 3D-E**). A similar finding was observed with cisplatin, with a 10.6-fold increase in IC50 observed following SIRT5 expression (**Fig. S4B**). To determine whether SIRT5 promotes chemoresistance through its enzymatic activity, we generated catalytic-deficient point mutants. Specifically, the critical interactions between Arg105, Tyr102, His158, and NAD in the acyl pocket of SIRT5 are essential for its enzymatic activity in desuccinylation and demalonylation (**Fig. 3F**) (18). We thus introduced specific point mutations at these residues that are known to significantly decrease SIRT5 catalytic activity, namely H158Y, R105M, and Y102F (18). Each of these point mutants demonstrated substantially attenuated effects on chemoresistance to both doxorubicin and cisplatin compared to WT SIRT5, despite expression at comparable levels (**Fig. 3D-E and Fig. S4B**). These results suggest that SIRT5-mediated chemoresistance is mediated through its enzymatic activity.

We then conducted a loss-of-function study using CRISPR knockout (KO) of SIRT5 in the SIRT5-expressing TNBC lines MDA-MB-231 and BT549 (See **Fig. 3C**). SIRT5 KO significantly enhanced chemosensitivity to doxorubicin and cisplatin compared to control cells that expressed scrambled gRNA (**Fig. 3G-J and Fig. S4C**). The chemosensitivity induced by loss of SIRT5 was limited, however, and we thus sought to explore potential functional redundancy with the other mitochondrial sirtuins, SIRT3 and SIRT4. Although SIRT4 is not expressed in breast cancer cells, we found that expression of SIRT5 in Hs578T cells slightly decreased SIRT3 protein levels (**Fig. S4D**), and conversely, SIRT5 KO in MDA-MB-231 cells led to an increase in SIRT3 protein expression. These observations suggest that SIRT3 may compensate in part for the absence of SIRT5 (**Fig. S4D**), and our subsequent studies below reveal how certain metabolites in the culture medium may contribute to this effect. Additionally, we found that SIRT5 expression is not correlated to chemosensitivity to the microtubule-destabilizing chemotherapy agent paclitaxel in TNBC cells (**Fig. S4E-G**). Overall, these findings establish a role of SIRT5 in mediating resistance to genotoxic chemotherapy agents, including doxorubicin and cisplatin, in TNBC.

### SIRT5 redirects glycolysis to the PPP to increase nucleotide pools

To investigate how SIRT5 alters metabolic programming leading to chemoresistance, we conducted Liquid Chromatography-Mass Spectrometry (LC/MS)-based metabolomics analyses on TNBC cell lines following SIRT5 overexpression (OE) or depletion compared to their respective isogenic controls. A total of 113 targeted metabolites were identified (**Table S7**). We found that SIRT5 OE in Hs578T cells significantly depleted the levels of glycolytic intermediates while increasing the levels of metabolic intermediates in the TCA cycle and nucleotide biosynthesis (**Fig. 4A-B**). Metabolite mapping underscored increases in NADPH and metabolic intermediates downstream of 6-phospho-D-gluconate within the PPP and nucleotide biosynthesis pathways, as well as increases in metabolic intermediates in glutaminolysis and the TCA cycle (**Fig. 4B**). As a control, we assessed metabolite profiles following expression of the three catalytic SIRT5 mutants. We found that compared to expression of wildtype SIRT5, expression of these mutants in Hs578T cells failed to increase the TCA intermediates, including citrate, α-ketoglutarate (α-KG), fumarate, and malate, as well as nucleotide biosynthesis intermediates such as NADPH, inosine monophosphate (IMP), guanosine monophosphate (GMP), and adenosine monophosphate (AMP) (**Fig. S5A-B**). Conversely, SIRT5 KO in MDA-MB-231 and BT549 TNBC cells resulted in increased glycolytic intermediates, while significantly decreasing nucleotide levels relative to cells expressing scrambled sgRNA control (**Fig. S5C-F**). Consistent with these findings, the metabolites involved in the TCA cycle, the PPP, and nucleotide biosynthesis are highly enriched in non-pCR TNBC primary tumors (See **Fig. 1C**). Collectively, our results suggested the hypothesis that SIRT5 catalytic activity redirects glucose metabolism from glycolysis towards the PPP, thereby promoting nucleotide biosynthesis, while at the same time increasing glutaminolysis to fuel the TCA cycle and promoting OXPHOS.

**Fig. 4.**
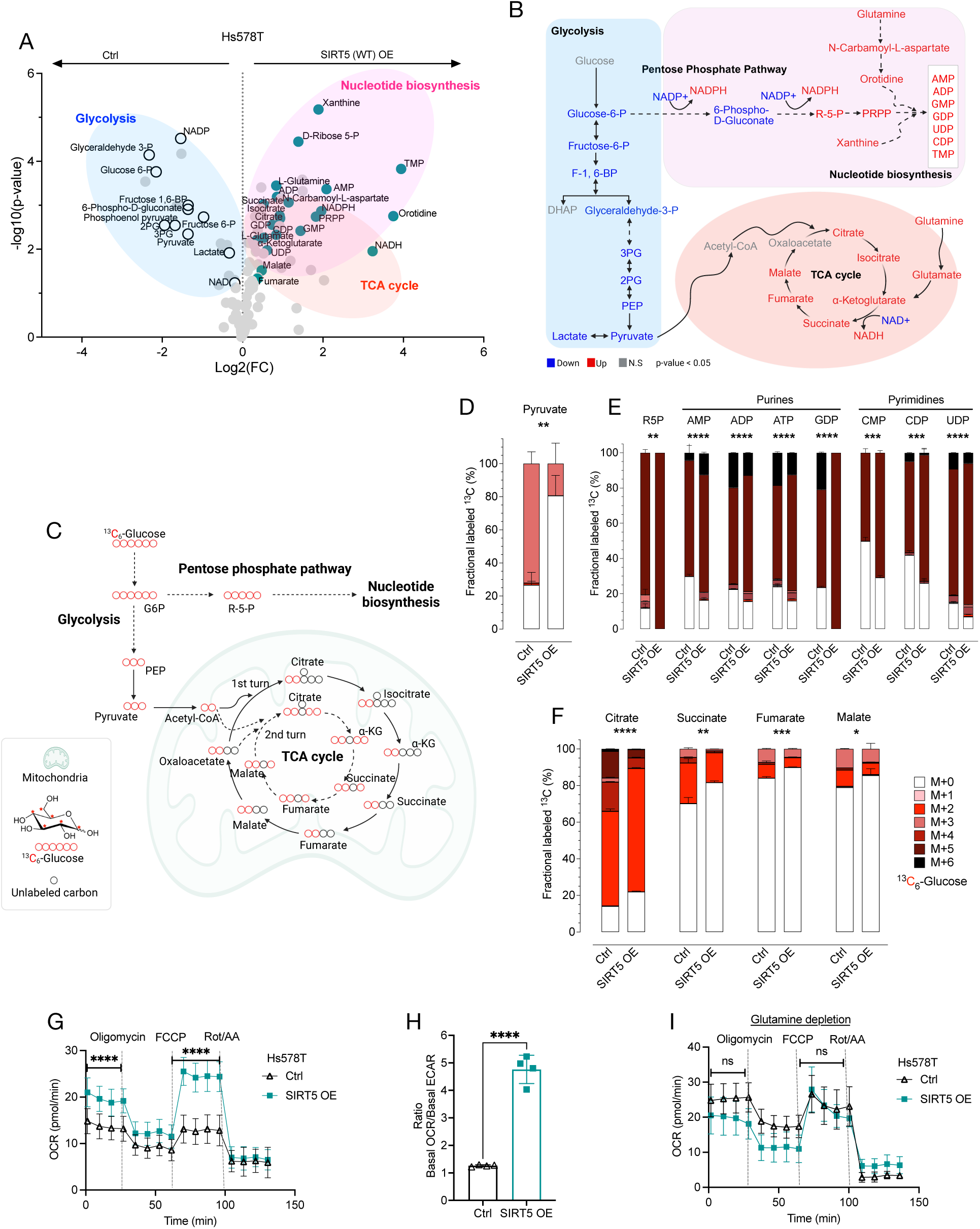
SIRT5 induces a metabolic switch from glycolysis toward the PPP for nucleotide biosynthesis. **A.** The volcano plot illustrates differentially expressed metabolites (n = 113) between Hs578T ctrl and SIRT5 expressing (OE) cells. Metabolites involved in glycolysis, nucleotide biosynthesis, and the TCA cycle are highlighted. **B.** Pathway mapping of metabolites profiled in panel A. Blue: decreased with SIRT5 OE; red: increased with SIRT5 OE. Metabolic pathways, including glycolysis, the TCA cycle, the PPP, and nucleotide biosynthesis, are included. **C.** Schematic representation of metabolic flux tracing by ^13^C-labeled glucose in glycolysis, the TCA cycle, and the PPP. **D-F**. Bar graphs display the percentage of fractionally labeled ^13^C detected in the indicated metabolic intermediates in glycolysis, the TCA cycle, and the PPP. **G**. Seahorse mitochondrial stress test shows an increase in oxygen consumption rate (OCR) in Hs578T/SIRT5 OE cells compared to control cells. **H**. Bar graphs present the ratio of basal OCR over basal ECAR in Hs578T/control and SIRT5 OE cells. **I.** Seahorse mitochondrial stress test shows elimination of increased OCR in Hs578T/SIRT OE cells cultured in glutamine-depleted media. ns indicates non-significance; Statistical significance is indicated with * p-value < 0.05; ** p-value < 0.01; *** p-value < 0.001; **** p-value < 0.0001.

To further explore this hypothesis, we conducted ^13^C-glucose metabolic flux tracing with a specific focus on metabolic interplay between glycolysis, the TCA cycle, the PPP, and nucleotide biosynthesis (**Fig. 4C**). Consistent with our metabolomics results, SIRT5 OE significantly diminished the fractional labeling of pyruvate from ^13^C-glucose, while markedly increasing the fractional labeling of ^13^C in R-5-P within the PPP as well as purine and pyrimidine nucleotides, compared to Hs578T control cells (**Fig. 4D-E**). Additionally, SIRT5 OE consistently resulted in a reduced fractional labeling of ^13^C from glucose in citrate, succinate, fumarate, and malate within the TCA cycle relative to the control Hs578T cells (**Fig. 4F**). Together, these results support the conclusion that elevated SIRT5 expression facilitates a metabolic switch from glycolysis to the PPP for nucleotide biosynthesis, leading to a diminished contribution of glycolysis to the TCA cycle.

These findings prompted us to look further into the TCA cycle and OXPHOS activity, given the increase in the TCA metabolites induced by SIRT5 (see **Fig. 4A-B**) despite a concomitant reduced contribution to their synthesis from glycolysis (**Fig. 4F**). We therefore conducted Seahorse assays to evaluate mitochondrial respiration and glycolysis in SIRT5 OE or depletion TNBC cells compared to their respective control. SIRT5 OE significantly enhanced the oxygen consumption rate (OCR) while decreasing the extracellular acidification rate (ECAR), a surrogate for glycolysis, in Hs578T cells, as indicated by both basal and maximal respiration rates, as well as the ratio of basal OCR over basal ECAR (**Fig. 4G-H**). In contrast, SIRT5 KO in MDA-MB-231 and BT549 cells significantly increased ECAR relative to their control cells (**Fig. S5G-H**). In alignment with the metabolomics analysis, the three SIRT5 mutants in Hs578T were significantly attenuated in their ability to affect OCR compared to wild-type SIRT5 (**Fig. S5I**).

Given the potential role of increased glutamine levels we observed in Hs578T SIRT5 OE cells to sustain the TCA and OXPHOS (See **Fig. 4A-B**), we then tested the dependence on glutamine for oxidative metabolism in this setting. We assessed OCR in glutamine-depleted media, comparing the effects of glutamine depletion in SIRT5 OE Hs578T cells versus control cells. Indeed, glutamine depletion eliminated the significant increase in OCR seen with SIRT5 OE (**Fig. 4I**). Together, these results suggest that SIRT5, through its enzymatic activity, increases OXPHOS in a glutamine-dependent manner, while at the same time diverting glycolysis to fuel the PPP and consequently reducing the contribution of glycolysis to the TCA anaplerosis.

### SIRT5 facilitates glutamine catabolism to sustain the TCA cycle and altered nucleotide pools that underlie chemoresistance

To test the role of glutamine in the setting of high SIRT5 levels, we conducted ^13^C-glutamine metabolic flux tracing with a focus on the flux between glutaminolysis, the TCA cycle, and nucleotide biosynthesis (**Fig. 5A**). The results demonstrated that SIRT5 increased the fractional labeling from ^13^C-glutamine to glutamate and multiple TCA metabolites, including α-KG, succinate, fumarate, and malate (**Fig. 5B-C**). The fractional ^13^C labeling of aspartate, as well as orotidine and UTP, were also significantly increased (**Fig. 5D-E**). Aspartate is a crucial precursor for pyrimidine nucleotide biosynthesis and is vital for cell proliferation and survival (43). These findings support the conclusion that SIRT5 promotes glutamine catabolism, providing a carbon source to sustain the TCA and nucleotide biosynthesis while mitigating the reliance on glucose-derived carbons to fuel the TCA cycle.

**Fig. 5.**
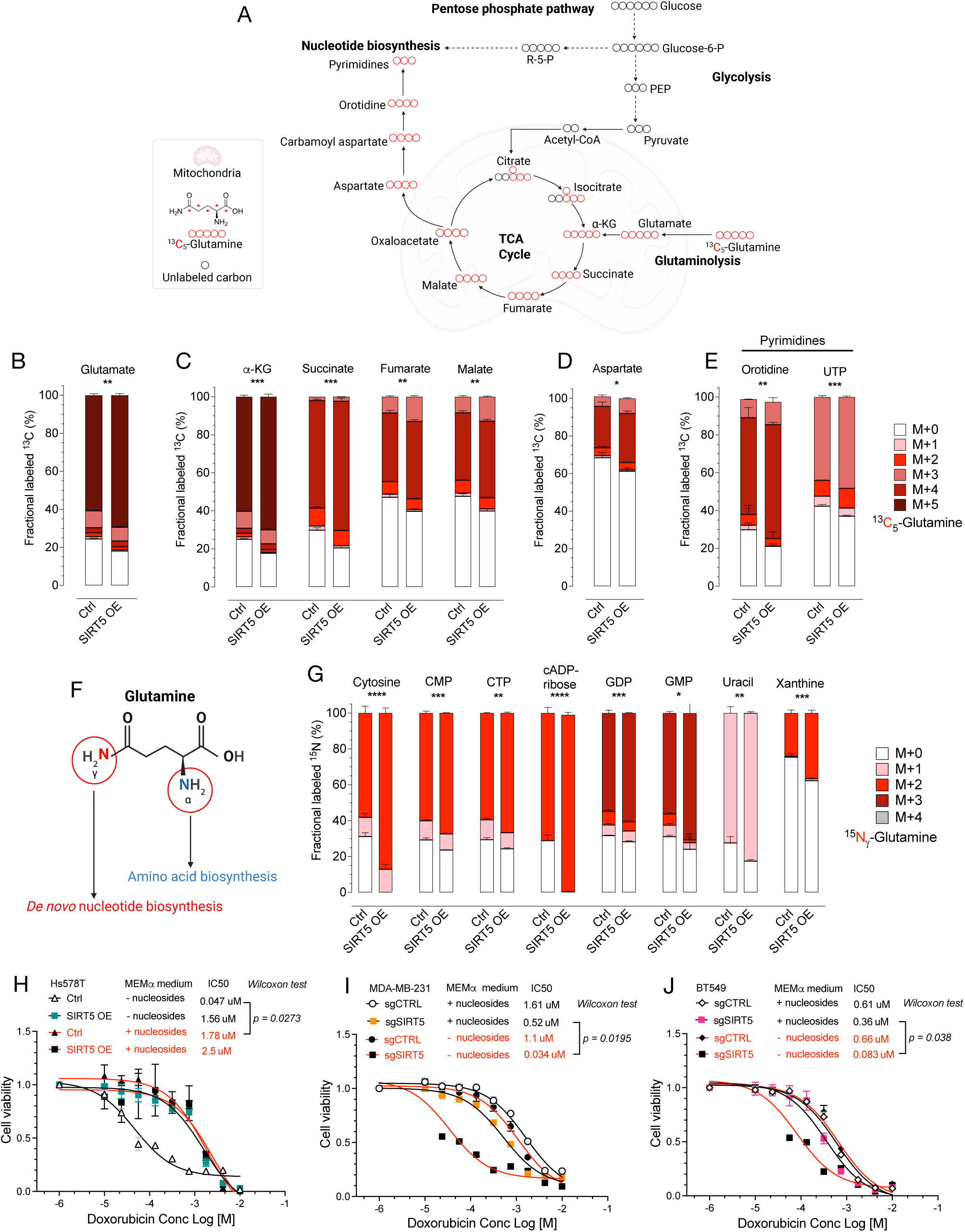
SIRT5 enhances glutamine metabolism to support the TCA cycle and nucleotide biosynthesis. **A.** Schematic representation of metabolic flux tracing of ^13^C-labeled glutamine in key metabolic pathways. **B-E**. Bar graphs illustrate the percentage of fractionally labeled ^13^C detected in glutamate, TCA intermediates, aspartate, and pyrimidines. **F**. Schematic diagram shows the nitrogen metabolic footprint of glutamine. Glutamine has two nitrogen atoms (α and γ) that can be donated, serving amino acid biosynthesis and de novo nucleotide biosynthesis, respectively. **G**. Bar graphs show the percentage of fractionally labeled ^15^N detected in indicated nucleotides and nitrogenous bases. **H-J**. Dose-response curves for Hs578T/SIRT5 OE cells versus control (H) and MDA-MB-231/SIRT5 KO and BT549/SIRT5 KO (J) cells versus their respective controls, following doxorubicin treatment for 72 hrs. Corresponding IC50 values are provided for cells grown in nucleoside-depleted or replenished MEMα culture media. Statistical significance is indicated as follows: *p < 0.05; **p < 0.01; ***p < 0.001; ****p < 0.0001.

Additionally, glutamine can serve as a crucial source of nitrogen for de novo nucleotide biosynthesis and the synthesis of amino acids (44). To establish whether SIRT5 regulates glutamine catabolism for nucleotide biosynthesis, we performed ^15^Nγ-glutamine isotope tracing (**Fig. 5F**). Consistently, SIRT5 significantly increased the fractional labeling from ^15^Nγ-glutamine to both purine and pyrimidine biosynthesis (**Fig. 5G**). Given that nucleotide pools are known to govern chemoresistance in many tumor settings, we postulated that SIRT5 regulates glucose and glutamine metabolism to induce chemoresistance through alteration of nucleotide pools (45).

To test this hypothesis, we assessed the effect of exogenous nucleoside supplementation, which can be salvaged to produce nucleotides, on doxorubicin chemosensitivity in TNBC cells under conditions of high or low SIRT5 levels. Indeed, nucleoside supplementation increased resistance of SIRT5-deficient Hs578T control cells to doxorubicin, enhancing the IC50 by 39 times—a response comparable to that observed in SIRT5 OE Hs578T cells in the absence of supplementation (**Fig. 5H**). Notably, nucleoside supplementation did not further increase resistance in SIRT5-overexpressing Hs578T cells (**Fig. 5H**). Correspondingly, nucleoside supplementation resulted in highly significant increases in doxorubicin resistance in SIRT5 KO MDA-MB-231 and BT549 cells, but had no effect on the respective control (scrambled sgRNA) cells, which express high endogenous levels of SIRT5 (**Fig. 5I-J**). Furthermore, SIRT5 OE significantly increased resistance to gemcitabine, a nucleoside analog that competitively inhibits nucleoside metabolism (**Fig. S6A**). Conversely, SIRT5 KO in MDA-MB-231 cells dramatically decreased resistance to gemcitabine (**Fig. S6B**). Collectively, these findings support the conclusion that SIRT5 plays a central role in enhancing nucleotide pools that govern chemosensitivity. Cells with high endogenous or ectopic SIRT5 exhibit chemoresistance that is not further increased by nucleoside supplementation, while cells with low SIRT5 levels can be rendered comparably chemoresistant by nucleoside supplementation.

### SIRT5 drives cellular dependence on glutamine as a bioenergetic substrate through activation of oncogenic MYC

Having identified nucleotide pools derived from enhanced glutamine metabolism as a key contributor to SIRT5-induced chemoresistance, we had yet to explain the increased glutamine levels observed following SIRT5 expression. Glutamine is imported into cells via solute carrier (SLC) transporters, specifically from the SLC1, SLC6, SLC7, SLC36, and SLC38 families (44). Notably, we found that SLC38A1 and SLC38A2, two key glutamine transporters, were significantly upregulated in SIRT5 OE Hs578T cells compared to controls (**Fig. S6C**). Conversely, these transporters were significantly downregulated in MDA-MB-231 cells with SIRT5 KO relative to control cells (**Fig. S6D**). MYC, which we previously linked to tumors with elevated SIRT5 (see **Fig. 2E-F and Fig. S3G**), transcriptionally regulates SLC transporters including the SLC38 genes, increasing the presence of SLC38A1 and A2 transporters in the cell membrane and thus enhancing glutamine uptake (46, 47). To further test the relationship between SIRT5 and MYC activity in human breast cancers, we analyzed an additional proteogenomic dataset obtained from the cBioPortal CPTAC database (48). GSEA of differentially expressed proteins between breast tumors with SIRT5 gain/amplification and those with diploid SIRT5 revealed that MYC targets consistently ranked among the top significantly enriched hallmark pathways in SIRT5 gain/amplified tumors at the protein level (**Fig. S6E**). Collectively, these results support the notion that SIRT5 promotes glutamine catabolism in part by regulating MYC activation.

SIRT5 has been reported to stabilize c-MYC through desuccinylation at the K369 site, promoting malignant progression in chordoma (49). We thus examined c-MYC protein levels in the presence and absence of SIRT5 in Hs578T, MDA-MB-231, and BT549 cells. As expected, c-MYC levels were elevated in SIRT5-overexpressing Hs578T cells, while they were reduced in SIRT5 KO cells in both MDA-MB-231 and BT549 cell lines compared to the controls (**Fig. S6F**). Next, we fractionated the cells and measured total succinylated lysine (Su-K) levels in both the cytosolic and mitochondrial compartments. Notably, SIRT5 OE resulted in a decrease in Su-K in Hs578T cells compared to the control, while SIRT5 KO led to an increase in Su-K in MDA-MB-231 cells, particularly within the mitochondria (**Fig. S6G-H**). We noted that the regions exhibiting significant changes in the Su-K cluster around 50 kDa, which corresponds to the molecular weight of the MYC protein (**Fig. S6G-H**). We then conducted immunoprecipitation (IP) followed by Western blot analysis of c-MYC to further investigate lysine succinylation. As anticipated, SIRT5 KO decreased MYC protein levels while simultaneously increasing Su-K on MYC (**Fig. S6I**). Thus, SIRT5 desuccinlyates MYC and increases its protein levels in breast cancer cells.

### SIRT5 redirects glycolysis to the PPP through facilitating the conversion of 6-phospho-D-gluconate to R-5-P by demalonylating 6-PGD

Our metabolite mapping suggests that SIRT5 may promote the conversion of 6-phospho-D-gluconate to R-5-P, thereby facilitating the PPP (see **Fig. 4B**). This conversion is primarily mediated by the key enzyme 6-phosphogluconate dehydrogenase (6-PGD), which catalyzes the oxidative decarboxylation of 6-phosphogluconate to R-5-P, generating NADPH and carbon dioxide (50). Notably, SIRT5 is unique in its localization to both the cytosol and mitochondria, while SIRT3 and SIRT4 are exclusively confined to the mitochondria (19). This dual localization implies that SIRT5 has the potential to regulate 6-PGD activity in the cytosol. To explore this possibility, we fractionated cells to determine whether SIRT5 influences lysine demalonylation in the cytosolic compartment. Indeed, SIRT5 OE decreased lysine malonylation in Hs578T cells compared to controls in the cytosol (**Fig. 6A**). In contrast, SIRT5 KO led to an increase in cytosolic lysine malonylation in MDA-MB-231 cells (**Fig. 6B**). Notably, a significant difference in lysine malonylation was observed around 50 kDa, which correlates with the molecular size of the 6-PGD protein (**Fig. 6A-B**). To further investigate the potential role of SIRT5 in the demalonylation of 6-PGD, we analyzed the lysine malonylome dataset from a prior analysis in *SIRT5^null^* and WT mice (22). This revealed a SIRT5-dependent malonylated lysine residue at position 59 in 6-PGD. (**Fig. 6C-D**). Moreover, murine 6-PGD Lys59 is conserved in humans (**Fig. 6E**).

**Fig. 6.**
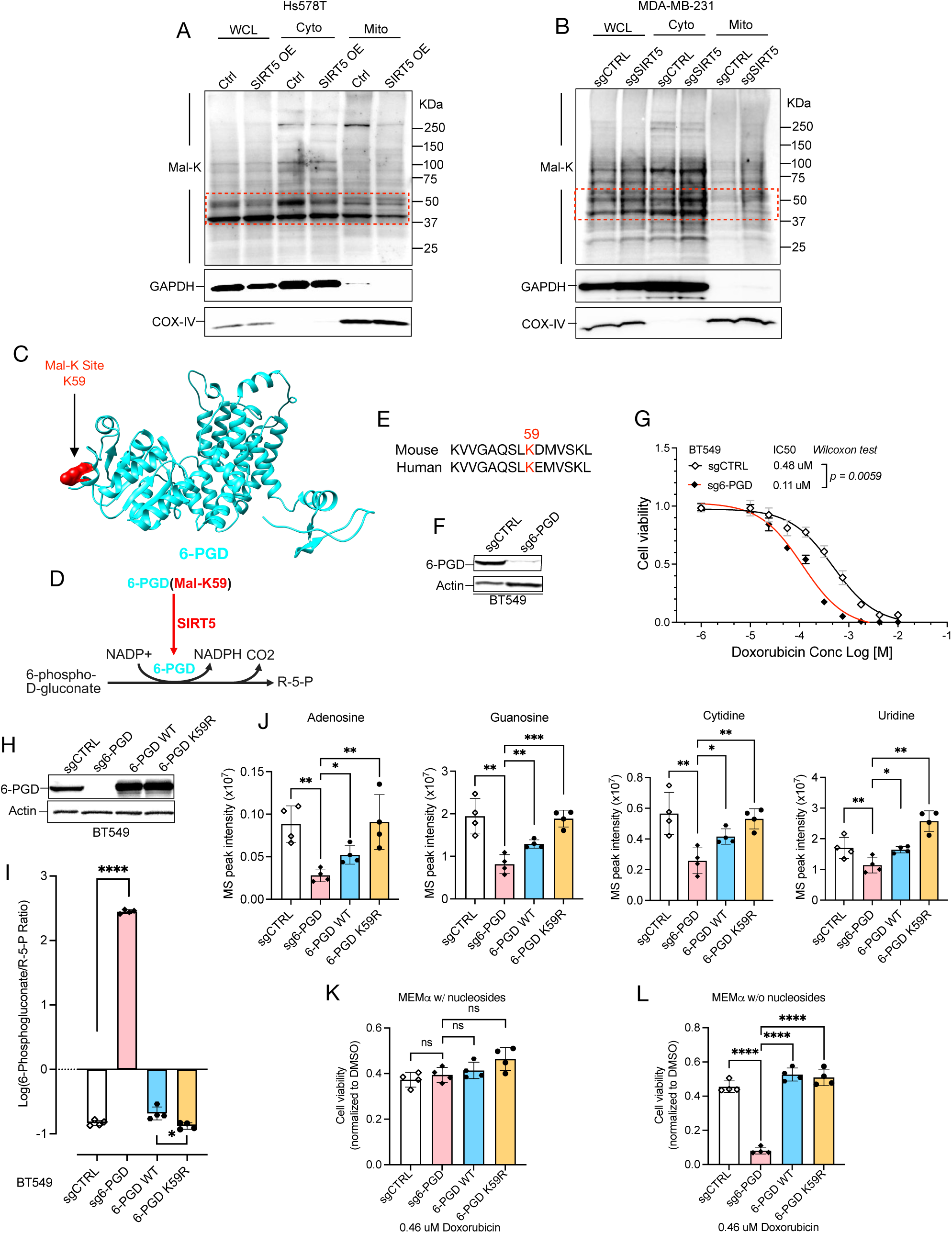
SIRT5 promotes the conversion of 6-phospho-D-gluconate to R-5-P for nucleotide biosynthesis by regulating 6-PGD demalonylation. **A-B**. Western blot showing levels of malonylated lysine (Mal-K) in whole cell lysate (WCL), cytosolic proteins (Cyto), and mitochondrial proteins (Mito) in Hs578T (A) and MDA-MB-231 (B) cells. Each panel includes quantification of Mal-K, with COX-IV and GAPDH used as loading controls. **C.** The crystal structure of 6-PGD (AlphaFold Q9DCD0) shows the Mal-K site at position 59. **D.** Schematic showing SIRT5 demalonylation of lysine residue on 6-PGD promotes conversion of 6-phospho-D-gluconate to R-5-P. **E.** 6-PGD^K59^ is conserved between mouse and human. **F.** Western blot shows 6-PGD levels in BT549 cells following expression of sg6-PGD or control. **G.** Dose-response curves show decreased sensitivity of BT549 cells to doxorubicin in the absence of 6-PGD; IC50 values are indicated. **H.** Western blot shows 6-PGD levels in BT549 cells expressing sgCTRL, sg6-PGD (KO), or reconstituted with 6-PGD WT or K59R mutant. **I.** Bar graphs show log10-transformed 6-Phosphogluconate/R-5-P ratio, highlighting the catalytic effects of 6-PGD on the conversion of 6-Phospho-D-gluconate to R-5-P. **J.** Bar graphs depict the mass spectrometry (MS) peak intensity of the indicated nucleosides in BT549 cells with or without 6-PGD WT and mutant reconstitution. **K-L**. Bar graphs show the drug sensitivity of BT549 cells expressing sgCTRL, sg6-PGD, 6-PGD WT, and 6-PGD K59R to 0.46 µM doxorubicin treatment after 72 hrs (normalized to DMSO), in MEMα culture media with (K) or without (L) nucleosides. ns indicates non-significance; statistical significance is indicated as follows: *p < 0.05; **p < 0.01; ***p < 0.001; ****p < 0.0001.

To determine whether SIRT5 regulates 6-PGD to enhance chemoresistance, we carried out CRISPR knockout of 6-PGD in the BT549 TNBC cells, which have high endogenous SIRT5 expression, and assessed sensitivity to doxorubicin (**Fig. 6F**). Indeed, 6-PGD KO significantly reduced the doxorubicin resistance compared to the control (**Fig. 6G**). To investigate whether SIRT5 promotes chemoresistance via demalonylation of the lysine residue K59 of 6-PGD, we rescued 6-PGD KO cells with either WT 6-PGD or a mutant form (K59R) that cannot be inhibited via malonylation, thus effectively mimicking SIRT5 demalonylation (**Fig. 6H**). Metabolomics analysis revealed that 6-PGD KO significantly decreased the conversion of 6-phospho-D-gluconate to R-5-P (**Fig. 6I**). Conversely, reintroduction of WT 6-PGD into 6-PGD KO BT549 cells restored this conversion, while the expression of the 6-PGD K59R mutant further enhanced conversion (**Fig. 6I**). Consequently, the 6-PGD KO led to a significant decrease in the levels of adenosine, guanosine, cytidine, and uridine. WT 6-PGD effectively rescued the levels of these nucleosides, with the K59R mutant further increasing their concentrations (**Fig. 6J**). Collectively, these results imply that PGD is activated and promotes nucleotide biosynthesis, leading to chemoresistance through the demalonylation of the K59 residue. To further elucidate whether 6-PGD mediates chemoresistance via altered nucleotide pools, we assessed the impact of exogenous nucleoside supplementation on doxorubicin sensitivity in these cells. Notably, exogenous nucleoside supplementation eliminated the differences in chemosensitivity observed between KO and rescue of 6-PGD (**Fig. 6K**). In contrast, depleting nucleosides from the culture media restored these differences (**Fig. 6L**). Overall, our data suggest that SIRT5 demalonylates 6-PGD at K59, facilitating the PPP and enhancing nucleotide pools, which in turn promotes chemoresistance.

### Elevated SIRT5 expression induces ATR replication stress checkpoint dependence in breast cancer

We hypothesized that metabolic rewiring and/or altered nucleotide pools in SIRT5-expressing tumors might be associated with potential vulnerabilities. To test this hypothesis, we utilized publicly available CRISPR dependency screening data from the DepMap database (DepMap Public 25Q3: depmap.org) to identify the genetic dependencies associated with SIRT5 expression across a diverse range of human breast cancer cell lines (51). This analysis revealed the replication stress checkpoint kinase ATR and its essential activating factors, TOPBP1 and RAD17, as the most significantly correlated common essential genes whose loss results in growth inhibition or cell death selectively in breast cancer cell lines with elevated SIRT5 expression (**Fig. 7A**). Importantly, this genetic dependency was only observed in breast cancer lines and was not evident in unselected tumor types (**Fig. 7B-C**). We thus hypothesized that altered nucleotide pools driven by SIRT5 might contribute to increased replication stress and the ATR checkpoint sensitivity. Supporting this hypothesis, existing evidence indicates that imbalanced nucleotide pools can trigger ATR-CHK1-dependent DNA replication stress signaling and DNA damage repair mechanisms, rendering cells dependent on the ATR checkpoint for proliferation (52, 53). In line with this idea, our metabolomics analysis indicated that SIRT5 OE selectively enhanced biosynthesis of particular nucleotides, including thymidine monophosphate (TMP) (**Fig. S7A**). Conversely, SIRT5 KO significantly decreased nucleotide biosynthesis, particularly in TMP (**Fig. S7B**). Supporting the link between enhanced pyrimidine biosynthesis and chemoresistance, previous studies have indicated that adaptive activation of de novo pyrimidine synthesis contributes to chemotherapy resistance in TNBC (12, 54-56).

**Fig. 7.**
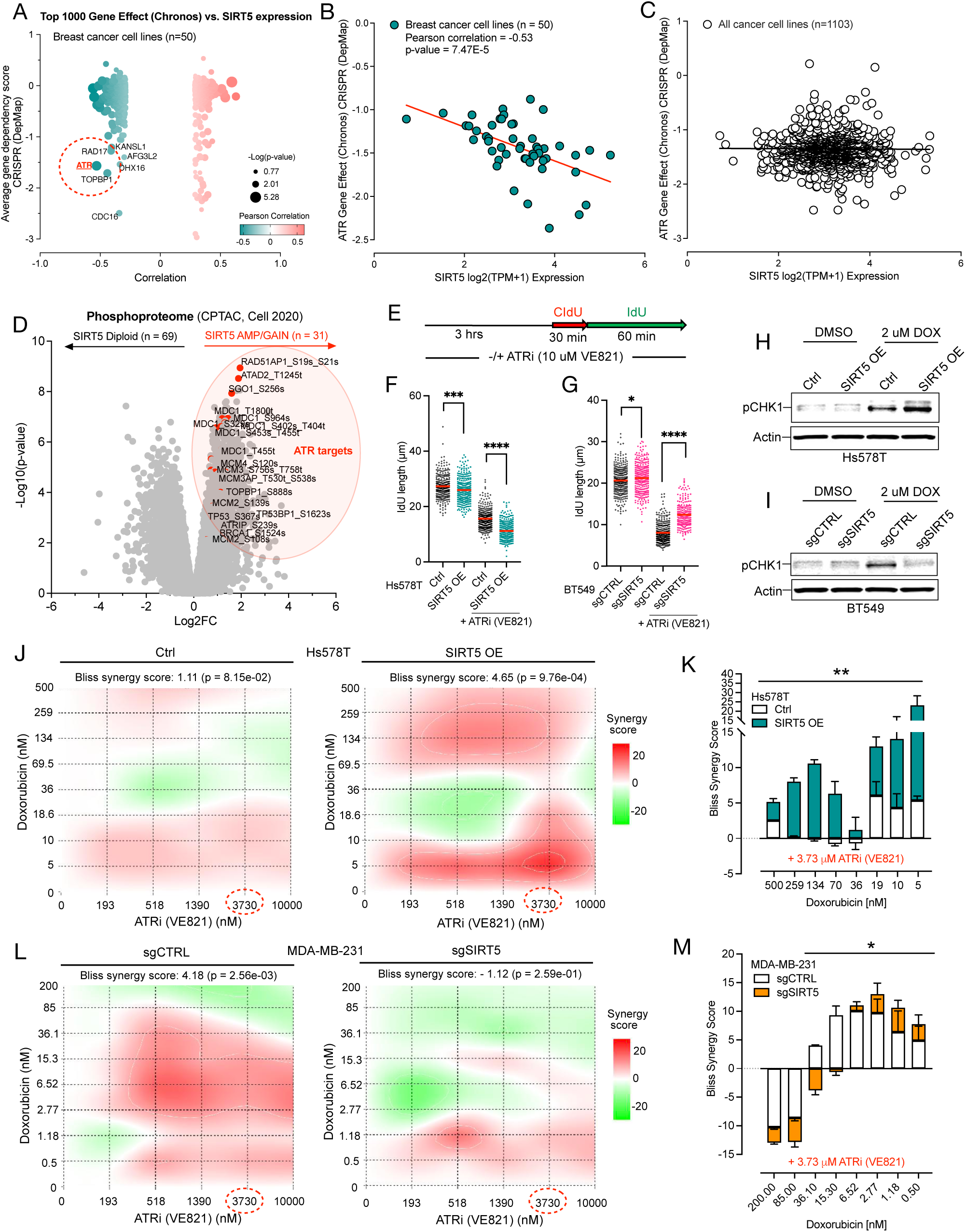
SIRT5 induces ATR checkpoint dependence as a key metabolic vulnerability to overcome SIRT5-induced chemoresistance in TNBC. **A.** Volcano plot shows results from the DepMap CRISPR screening demonstrating association of SIRT5 expression with dependency on ATR checkpoint factors, including ATR, TOPBP1, and RAD17, across 50 breast cancer cell lines. **B.** Dot plot highlights the correlation between ATR CRISPR dependency scores and SIRT5 expression across the 50 breast cancer cell lines, normalized within the Chronos matrix. **C.** The dot plot shows the correlation between ATR CRISPR dependency scores and SIRT5 expression across a broader dataset of 1103 cancer cell lines, also normalized in the Chronos matrix. **D.** Volcano plot shows the differentially expressed phosphoproteins in primary breast tumors harboring SIRT5 amplification/gain (n = 69) versus SIRT5 diploidy (n = 31). Significantly differentially expressed ATR target phosphoproteins are indicated in red. **E.** Schematic illustrates the DNA fiber assay measuring replication tracts. Cells are first incubated in 10 µM ATRi (VE821) or vehicle control for 3 hrs, then sequentially labeled with the thymidine analogs CldU for 30 minutes, followed by IdU for 60 minutes. **F-G**. The dot plots show the quantification of DNA replication tracts, measured by IdU length, in Hs578T and BT549 cells with or without SIRT5 expression. **H-I**. Western blots show the expression of pCHK1 in Hs578T and BT549 cells in the presence or absence of SIRT5 expression. Cells were treated with 2 µM doxorubicin or vehicle (DMSO) overnight. **J-M**. The synergistic interaction between doxorubicin and ATRi was determined using Synergy Finder, which calculates the Bliss synergy score in Hs578T cells (J) and MDA-MB-231 (L) cells in the presence or absence of SIRT5 expression. The Bliss synergy scores are quantified from three independent experiments for ARTi at a dosage of 3.73 µM and doxorubicin at different dosages for 5 days; results are shown in (K) and (M). Statistical significance is indicated as follows: *p < 0.05; **p < 0.01; ***p < 0.001; ****p < 0.0001.

To further elucidate the link between SIRT5 and replication stress signaling in human breast cancer, we conducted a differential expression analysis of the breast cancer phosphoproteomic dataset from the cBioPortal CPTAC database (**Fig. 7D**) (48). This analysis revealed that RAD51AP1, a direct target of the ATR kinase and a key player in homologous recombination DNA repair, was the most enriched phosphoprotein in breast tumors exhibiting SIRT5 gain/amplification (**Fig. 7D**) (57). Other highly differentially phosphorylated proteins included MDC1, which is not a direct ATR phosphorylation target but functions alongside ATR in replication stress response and DNA repair (**Fig. 7D**). Additionally, key ATR substrates including ATAD2, SGO1, MCM, TOPBP1, TP53, TP53BP1, ATRIP, and BRCA1 were found to be significantly upregulated in SIRT5 gain/amplified breast tumors (**Fig. 7D**). Importantly, these associations are consistent with the idea that elevated SIRT5 may be positively selected during tumor evolution as an adaptive mechanism to cope with rising replication-associated genome instability, thereby enabling continued proliferation under oncogenic and metabolic stress (58). Taken together, our findings suggest that elevated SIRT5 expands and modifies nucleotide pools that may promote DNA repair and chemoresistance, but at the expense of increased replication stress and dependence on the ATR checkpoint.

We next assessed directly the role of SIRT5 in ATR checkpoint dependence, carrying out DNA fiber assays to measure DNA replication fork progression (**Fig. 7E**). SIRT5 OE resulted in modest but significant attenuation of fork progression in Hs578T cells at baseline, and addition of the ATR inhibitor VE821 slowed fork progression to a much larger degree with SIRT5 OE compared to control cells (**Fig. 7F**). Conversely, in BT549 cells with high endogenous SIRT5 levels, SIRT5 knockout (SIRT5 KO) modestly accelerated fork progression, and ATR inhibition slowed fork progression significantly less in SIRT5 KO cells than in matched controls (**Fig. 7G**). We then tested how SIRT5 affected ATR checkpoint activation in response to genotoxic stress. Doxorubicin treatment in SIRT5 OE Hs578T cells resulted in a significant increase in phosphorylated levels of the key ATR substrate CHK1 compared to control cells (**Fig. 7H**). By contrast, BT549 cells with endogenous SIRT5 KO exhibited substantially lower pCHK1 levels upon doxorubicin treatment compared to controls (**Fig. 7I**). Thus, high levels of SIRT5 found in chemoresistant breast cancer confer ATR dependence and hyper-activation in response to DNA damage.

These findings led us to test the potential of targeting ATR as a critical metabolic vulnerability to counteract SIRT5-induced chemoresistance in TNBC. We carried out quantitative dose-response curves for chemotherapy in combination with the ATR inhibitor VE821. Significant synergy was observed between VE821 and doxorubicin in SIRT5 OE but not control Hs578T cells at inhibitor concentrations achievable in vivo (3uM) (**Fig. 7J-K**) (59). Furthermore, in chemoresistant MDA-MB-231 cells that express high endogenous SIRT5 levels, synergy was observed between VE821 and doxorubicin at baseline, but was abolished with SIRT5 KO (**Fig. 7L-M**). Collectively, these results suggest that targeting ATR represents a novel metabolic vulnerability to counteract SIRT5-induced chemoresistance in TNBC.

## Discussion

Prior studies have suggested that SIRT5 may play a potential oncogenic role in multiple cancers, including colorectal cancer (58, 60), ovarian cancer (61), melanoma (62), acute myeloid leukemia (63), and breast cancer (64). Here, we analyzed meticulously collected pretreatment TNBC tumor samples as a primary discovery approach, revealing SIRT5 overexpression as both a hallmark and a mediator of chemoresistance in this context. In keeping with this conclusion, we confirmed that SIRT5 frequently exhibits elevated expression in TNBC tumors, driven in part by genomic gain/amplification events. Our studies have uncovered in detail how high SIRT5 levels reprogram metabolism to support chemoresistance. Most importantly, we discovered an unanticipated liability associated with SIRT5 metabolic reprogramming, thus providing new and actionable insight into how metabolic adaptations that drive cancer progression and drug resistance may be exploited therapeutically.

Our findings point to SIRT5 as the nexus of a coordinated reprogramming of metabolism that underlies chemoresistance. Through enzymatic modulation of key substrates, SIRT5 diverts glycolytic metabolism to enhance nucleotide biosynthesis via the oxidative PPP, while promoting glutaminolysis to fuel OXPHOS and mitochondrial metabolism. The mechanisms of SIRT5 revealed through our TNBC models are strongly supported by our findings in primary tumor specimens. Analysis of both metabolite and proteome datasets we generated from primary breast cancer samples reveals a significant correlation between increased SIRT5 expression and the emergence of a highly oxidative phosphorylation (OXPHOS)-dependent energetic state within tumor cells. In addition to metabolic changes, our analysis of phosphoproteome data from primary tumors demonstrates that elevated SIRT5 levels are linked to modifications in DNA damage response and repair mechanisms, and particularly to activation of the ATR-dependent replication stress checkpoint. These alterations can contribute to the development of resistance to DNA damage-inducing therapies, but they also underlie a potential vulnerability in these tumors that could be exploited therapeutically. Importantly, the concordance of proteomic and metabolomic analyses of patient-derived tumor specimens we analyzed underscores the physiological relevance of these findings. Clinically, high SIRT5 levels correlate with worse relapse-free survival in chemotherapy-treated TNBC patients – but not in patients who did not receive chemotherapy - indicating its role as both a biomarker and mediator of chemoresistance in these tumors.

Further functional studies revealed how SIRT5 orchestrates the chemoresistant phenotype in TNBC through its enzymatic activity. We find that SIRT5 facilitates a metabolic switch in glucose metabolism from glycolysis to the PPP through demalonylation of 6-PGD, thus augmenting the conversion of 6-phospho-D-gluconate to R-5-P within the PPP and ultimately enhancing nucleotide biosynthesis. We demonstrate that manipulation of nucleotide pools can quantitatively recapitulate effects of SIRT5 on chemosensitivity, providing direct evidence for altered nucleotide metabolism as a mechanism of SIRT5-induced chemoresistance.

In parallel, SIRT5 supports the energetic demands of rapidly proliferating tumors by augmenting glutamine levels and glutaminolysis. Our data, alongside findings from other studies, indicate that SIRT5 promotes glutamine catabolism through MYC de-succinylation and activation that support the TCA cycle and nucleotide biosynthesis (49, 65). This interpretation is consistent with our findings that SIRT5 increases the contribution of glutamine-derived carbons and nitrogen to TCA substrates and nucleotides, respectively. This compensatory mechanism safeguards against carbon loss due to glycolytic diversion, thereby sustaining the energetic demands of TNBC cells during chemotherapy. Thus, the metabolic environment cultivated by SIRT5 expression enables TNBC cells to sustain their proliferative capabilities while simultaneously supporting robust repair to mitigate DNA damage under chemotherapeutic stress.

While SIRT5 has been linked to chemotherapy resistance in other contexts, our study is, to our knowledge, the first to demonstrate an actionable therapeutic vulnerability linked to SIRT5 metabolic reprogramming. We find that the top genetic dependency associated with SIRT5 expression in breast cancer centers on the ATR replication stress checkpoint, which maintains genome stability by stabilizing replication forks and promoting DNA repair (66). SIRT5 genomic gain/amplification events are associated with increased phosphorylated ATR substrates in primary breast cancer samples, including proteins involved in DNA repair and replication. In addition, overexpression of SIRT5 in our TNBC models is sufficient to slow replication fork progression, and ATR inhibition induces a significantly greater effect on fork progression in cells with high SIRT5 levels. This ATR checkpoint dependence may relate to replication stress from imbalanced nucleotide pools, as both loss and gain of SIRT5 expression preferentially change TMP levels (see Fig. S7). Increased TMP induced by SIRT5 is particularly notable given several recent studies demonstrating selective dependence of TNBC tumors on high pyrimidine levels (12, 54-56). Collectively, our findings unveil an intricate interplay between metabolic state and energetic demands, nucleotide biosynthesis, replication stress, and DNA repair mechanisms that, on the one hand, drive tumor progression and chemoresistance, and on the other create an exploitable susceptibility. Indeed, we find that ATR inhibition not only decreases chemosensitivity in TNBC, but does so synergistically in the setting of elevated SIRT5. Thus, targeting ATR to reverse chemoresistance in SIRT5-overexpressing TNBC provides both a disease context and a tumor-specific biomarker for this approach. These discoveries address both tumor selectivity and therapeutic index, which are the major ongoing challenges to the clinical implementation of ATR inhibitors for cancer therapy (67).

In conclusion, our findings establish SIRT5 as a critical mediator of metabolic reprogramming and chemoresistance in TNBC. These results contribute to an evolving narrative regarding the role of metabolic adaptation in cancer survival and associated tumor-specific vulnerabilities. We present a compelling case for further investigation of SIRT5 as a predictive biomarker for chemoresistance in TNBC, and we propose that targeting ATR represents a novel strategy for overcoming SIRT5-mediated chemoresistance in this challenging subtype of breast cancer.

## Acknowledgments

This work was supported by grants from the ESSCO MGH Breast Cancer Research Fund (to L.M. Spring), Ludwig Center at Harvard funding (to L.W. Ellisen and M.C. Haigis), and the Tracey Davis Memorial Breast Cancer Research Fund (to L.W. Ellisen). Additionally, Z. Ren is supported by the American Association for Cancer Research (AACR) Breast Cancer Fellowship. We thank Dr. Brian N. Dontchos for his generous assistance with tumor biopsy collection, and Agustina Maccio and Preshita Dave for their help with bioinformatics for this project. We extend our gratitude to all the patients and investigators who participated in the biopsy project.

## Author contributions

**A. Z. Ren**: Conceptualization, investigation, methodology, formal analysis, data curation, software, data visualization, writing original draft, writing-review and editing; **T. Bernasocchi, K. Kurmi, B. Ordway, and R. Morris**: Data curation, software, formal analysis, data visualization, writing-review and editing; **T. Bernasocchi, C.X. Guo, K. Jiang, S. Joshi, E. Zaniewski, X. Li, and K.N. Islam**: Methodology, investigation, validation, writing-review and editing; **G.X. Wang, and S-H. S. Chou:** Sample acquisition and writing, review, and editing. **L.M. Spring and V. I. Bossuyt**: Sample acquisition, data curation, writing-review, and editing. **G. Lam, M. Lawrance, and E. Rheinbay**: Data analysis, data curation, software, data visualization, writing-review, and editing. **I. Sanidas, L. Zou, W. Haas, R. Mostoslavsky, and M.C. Haigis**: Resources, investigation, and writing-review and edit. **L.W. Ellisen**: Conceptualization, resources, formal analysis, supervision, funding acquisition, writing original draft, writing review, and editing.

## Data availability

The datasets downloaded from TCGA, METABRIC, and cBioPortal are analyzed and the coding scripts are uploaded to a GitHub repository for this project under rheinbay_lab (https://github.com/rheinbaylab/Ellisen_Lam_SIRT5). The TMT-proteomics raw data reported in this paper are available on MASSIVE under the accession number MSV000101354 (ftp://MSV000101354@massive-ftp.ucsd.edu).

## Materials and Methods

### Patient specimen collection

Biopsy samples were collected from standard ultrasound-guided core needle breast biopsies at the time of initial tissue diagnosis at Massachusetts General Hospital, with protocol approved by the Massachusetts General Brigham Institutional Review Board (001865), and with informed consent obtained from all patients. After breast tissues were biopsied, they were snap-frozen immediately on dry ice and later stored in a freezer at -80 ^0^C until research (24). All consented patients had their clinical data collected and entered into a secure REDCap (Research Electronic Data Capture) database. Medical records of these patients were periodically reviewed to collect data on demographics, biopsy results, tumor and treatment characteristics, and long-term outcomes.

### Cell lines

The human breast cancer cell lines, including Hs578T, MDA-MB-231, and BT549, are obtained from ATCC. These human breast cancer cell lines were cultured in RPMI medium (Gibco) supplemented with 10% heat-inactivated fetal bovine serum (FBS) and 1% Pen-Strep (Gibco) at 37 °C with 5% CO2. All cell lines are regularly tested for the absence of Mycoplasma.

### Constructs

Human SIRT5 lentiviral vector pLV[Exp]-Puro-CMV>hSIRT5[NM_001376804.1] and SIRT5 mutation lentiviral vectors, including pLV[Exp]-Puro-CMV>{hSIRT5*(Y102F)}, pLV[Exp]-Puro-CMV>{hSIRT5*(R105M)}, and pLV[Exp]-Puro-CMV>{hSIRT5_H158Y*}, were obtained from VectorBuilder. Human 6-PGD lentiviral vector with PAM domain mutation pLV[Exp]-Bsd-EF1A>{hPGD[NM_002631.4]*_mut_PAM} and 6-PGD mutant lentiviral vector with PAM domain mutation pLV[Exp]-Bsd-EF1A>{hPGD[NM_002631.4]*(K59R)_mut_PAM} were purchased from Vector Builder. Lentiviral particles produced from the obtained plasmids were used for the expression of their protein in the Hs578T, MDA-MB-231, and BT549 cell lines.

### Lentivirus production

Lentiviral particles were generated by transient co-transfection of 293T cells with a lentivirus-based vector expressing either the full-length clone of the desired gene or a CRISPR all-in-one lentiviral vector to target the specific gene in the genome. Briefly, a 100-mm dish seeded with 3 × 10^^6^ cells was transfected with 20 µg of DNA plasmids that include 10 µg of expression vector and 10 µg of lentiviral packing mix (Cat#. A43237) diluted in 500 µL OPTIMEM (Solution A). Lipofectamine 2000 (Cat#. 11668019) was used for transient transfection in 500 µL OPTIMEM (Solution B). Solutions A and B were mixed and incubated at room temperature for 20 minutes to form the transfection complex, which was then added to 293T cells. After 48 hours of transfection, the supernatant was collected, centrifuged at 2,000 rpm for 10 minutes, and filter-sterilized.

### CRISPR lentiviral transduction

Mammalian CRISPR lentiviral vector (pLenti-U6-sgRNA-SFFV-Cas9-2A-Puro) against human SIRT5 with gRNA sequences (target 1: CGACTTGGGACAATCTGGAG, target 2: CAGGCAAATCTGGTTTCGTG, and target 3: AGTGGTGTTCCGACCTTCAG) was purchased from ABM. Mammalian CRISPR lentiviral vector against human 6-PGD with gRNA sequences (target 1: GACGGGTACGCCGTATTCCA and target 2: TGACGGGTACGCCGTATTCC) was obtained from VectorBuilder. CRISPR lentiviral vectors against human SIRT5 and 6-PGD, and a CRISPR control containing a non-targeting scramble gRNA lentiviral vector were packaged in 293T cells and infected into cells. Briefly, MDA-MB-231, BT549, or Hs578T cells were seeded at a concentration of 1 × 10^^5^ cells per 12-well plate for 24 hours, treated with 250 µl of viral solution containing 10 µg/ml polybrene for 1 hour, and incubated in RPMI/10% FBS without antibiotics for 24 hours. Cells were then expanded into a 10-cm dish with selective antibiotics.

### Antibodies

Antibodies against SIRT3 (Cat. # 5490S, 1:1000), SIRT4 (Cat. # 69786S, 1:1000), SIRT5 (Cat. # 8782S, 1:1000), pCHK1 (Cat. # 2348T, 1:1000), and Mal-K (Cat. # 14942S, 1:1000) were obtained from Cell Signaling Technology. Actin antibody (Cat. # A2066, 100 µL, 1:5000) was purchased from Sigma. Antibody against c-MYC (Cat. # ab32072) for WB (1:1000) and IP was obtained from Abcam. Su-K (Cat. # PTM-419, 1:1000) was obtained from PTM Bio. GAPDH (Cat. # GTX627408, 1:5000) was from GeneTex. 6-PGD antibody (Cat. # 14718-1-AP, 1:1000), COX-IV (Cat. # 112421AP, 1:1000), and rabbit IgG control antibody (Cat. # 3000-0-AP) were purchased from Proteintech.

### Immunoblotting

Cells were extracted in RIPA solubilization buffer (50 mM Tris-HCl, pH 7.5, 150 mM NaCl, 0.5mM MgCl2, 0.2mM EGTA, 1% Triton X-100), including protease and phosphatase inhibitors (A32965). Twenty micrograms of protein were loaded onto 7-12% SDS-polyacrylamide gels and transferred to Immobilon membranes. Blots were probed overnight at 4°C with the indicated antibodies and developed using chemiluminescence (Thermo Fisher Scientific).

### Crosslink immunoprecipitation

Immunoprecipitation was performed using a protocol adapted from the Pierce Crosslink Magnetic IP/Co-IP Kit (88805). Before covalent crosslinking, protein A/G magnetic beads were pre-washed twice with 1x modified coupling buffer. The antibody was then coupled to the beads at room temperature (RT) for 30 minutes on a rotator. Following this, the prepared beads were washed three times with 1x modified coupling buffer and crosslinked to the antibody using disuccinimidyl suberate (DSS) for 1 hour at RT on a rotator. The crosslinked beads were subsequently washed three times with Elution Buffer, followed by twice with IP Lysis/Wash Buffer. 500 µg of cell lysates were incubated with 25 µL of crosslinked beads overnight at 4 °C on a rotator. After incubation, the unbound fraction was collected, and the beads were washed twice with IP Lysis/Wash Buffer and once with ultrapure water. Bound antigens were then eluted for downstream analysis.

### Immunohistochemistry

Formalin-fixed and paraffin-embedded patient tissues were sectioned to a thickness of 5 µm, followed by deparaffinization in xylene and rehydration through a series of decreasing ethanol-H2O solutions. Antigen retrieval was performed by heating the sections to 100 °C for 30 minutes in 0.01 M trisodium citrate buffer at pH 6.0. Endogenous peroxidase activity was inhibited by incubating the slides in 3% hydrogen peroxide for 10 minutes. Subsequently, the tissues were blocked using Dako Serum-Free Protein Block and incubated with primary antibodies in Invitrogen antibody diluent containing 1% BSA at RT for 1 hour or overnight at 4 °C. After washing, the tissues were incubated with Dako Envision Labeled Polymer-HRP anti-mouse or anti-rabbit secondary antibodies at room temperature for 30 minutes. Following another wash, the samples were developed using DAB substrate (Cell Marque, Cat. 957D), after which sections were washed and counterstained with Harris Hematoxylin (Fisher Scientific). The sections were then dehydrated, mounted, and validated by a breast pathologist. The quantification of SIRT5 expression was performed on the entire view of IHC staining of non-pCR and pCR TNBC tumors. The expression levels were scored based on staining intensity and tumor cell area, using a weighted histoscore calculated as follows: (1 × 3% weak staining) + (2 × 3% moderate staining) + (3 × 3% strong staining).

### Drug dose response curve

A total of 2 × 10^5 cells were diluted to 40 cells/µL in 5 mL of standard RPMI medium supplemented with 10% FBS. For nucleoside supplementation and depletion experiments, the cells were diluted in nucleoside-depleting or replenishing MEMα culture media (Cat. #12561056 and Cat. #12571063) supplemented with 10% dialyzed FBS (GIBCO). From here, 2,000 cells in 50 µL of the prepared medium were seeded in triplicate across ten wells of a 384-well plate to allow for overnight cell adhesion. On the following day, doxorubicin and cisplatin were reconstituted to a stock concentration of 10 mM and loaded into a drug printer (Tecan D300e) for distribution. Doxorubicin was dispensed in a concentration range from 10 µM to 0 µM, while cisplatin was distributed in a range from 25 µM to 0 µM. After administering the drugs, the cells in the plate were centrifuged at 500 rpm to ensure the drugs precipitated in the medium. Subsequently, the cells were incubated with the drugs for 72 hours. The CellTiter-Glo assay (Cat. # G7571) was employed to measure cell viability and calculate the IC50 values. A total of 20 µL of the CellTiter-Glo solution was added to the experimental wells containing the cell culture medium. Control wells containing medium without cells were used to determine the background luminescence. The plate was then incubated for 10 minutes on an orbital shaker at RT to induce cell lysis and stabilize the luminescent signal in the dark. Luminescence readings were recorded with a plate reader, and the data were analyzed in Prism 10 to calculate IC50 values.

### Synergy score calculation

To assess the potential synergistic effects between ATR inhibitors (ATRi) and doxorubicin, we cultured 1,000 cells per well in standard RPMI medium supplemented with 10% FBS in 96-well plates. After 24 hours of incubation, the cells were treated with the respective drugs. The CellTiter-Glo (Cat. # G7571) assays were performed five days post-treatment to evaluate cell viability. A total of 50 µL of CellTiter-Glo solution was added to the experimental wells containing 100 µL of cell culture medium. The plate was then incubated for 10 minutes on an orbital shaker at RT to induce cell lysis and stabilize the luminescent signal in the dark. Luminescence readings were recorded with EnVision Nexus (revvity) plate reader, and the data were analyzed in Prism 10 to calculate IC50 values. The effects of the drug combination were analyzed relative to the growth of untreated cells. Bliss synergy scores were computed using the SynergyFinder Plus web application (68).

### DNA fiber assay

A total of 0.1 × 10^6 cells were seeded in twelve-well plates. The following day, for ATR inhibition, VE821 (10 µM) was added 3 hours prior to and maintained throughout the labeling process. Cells were labeled with 50 µM CldU (Sigma C6891-100MG) for 30 minutes, washed three times with equilibrated culture media, and subsequently labeled with 250 µM IdU (Sigma I7125-5G) for 60 minutes. After washing twice with 1× phosphate-buffered saline (PBS), the cells were scraped on ice in 500 µl of chilled PBS and centrifuged at 7,000 rpm for 6 minutes at 4°C, leaving approximately 100 µl of PBS for resuspension. Two 3 µl drops of cell suspension were placed near the frosted end of glass slides and air-dried for 2 minutes. Lysis buffer [200 mM Tris (pH 7), 50 mM EDTA (pH 8), 0.5% SDS] (8 µl per drop) was then added, and the mixture was incubated for 10 minutes at room temperature. The slides were tilted at 15° to allow DNA spreading, air-dried for an additional 10 minutes, fixed in a 3:1 methanol: acetic acid solution for 10 minutes, and washed three times with PBS. DNA was denatured in 2.5 N HCl for 90 minutes at room temperature, followed by washing in PBS and blocking in fiber-blocking buffer (2% BSA, 0.1% Tween 20 in PBS) for 30 minutes at 37°C. The slides were incubated with rat anti-BrdU (1:50; Abcam ab6326, CldU) and mouse anti-BrdU (1:250; BD 347580, IdU) for 60 minutes at 37°C, washed, and then incubated with Alexa Fluor 594 anti-rat IgG and Alexa Fluor 488 anti-mouse IgG (1:100; Invitrogen A-11001) for another 60 minutes at 37°C. Following washing, the slides were air-dried, mounted with ProLong Gold Antifade, and imaged on a Nikon Eclipse Ti2 microscope using a 40× air objective. Images were analyzed using Fiji.

### Seahorse assay

To measure oxygen consumption rate (OCR), cells were suspended in normal growth medium and seeded at 20,000 cells/well in Seahorse 96-well plates 24 hrs prior to the assay. Mitochondrial stress test was carried out in an XF96 Seahorse Analyzer (Seahorse Bioscience, Billerica, MA, USA). On the day of the assay, medium was replaced with pre-warmed (37 °C) 180 µl Seahorse XF medium (2 mM glutamine, 10 mM glucose, 1 mM pyruvate) and the plate was incubated for 1 hour at 37 °C in a non-CO2 incubator. Basal OCR measurements were recorded 4 times (mix: 3 min; wait: 2 min; measure: 3 min), followed by sequential injection of 1 µM oligomycin, 1 µM FCCP, and 0.5µM rotenone/antimycin with 4 readings (mix: 3 min; wait: 2 min; measure: 3 min) after each injection. To measure extracellular acidification rate (ECAR), cells were suspended in normal growth medium and seeded at 20,000 cells/well in Seahorse 96-well plates 24 hrs prior to the assay. Glycolysis stress assay was carried out in an XF96 Seahorse Analyzer (Seahorse Bioscience, Billerica, MA, USA). Before assay, the medium was replaced with pre-warmed (37 °C) 180 µl of XF Base Medium (Agilent Technologies) supplemented with 2 mM glutamine, and the mixture was incubated for 1 hour at 37 °C in a non-CO_2_ incubator. Baseline ECAR measurements were recorded 4 times (mix: 3 min; wait: 2 min; measure: 3 min), followed by sequential injection of 20 mM glucose, 1 µM oligomycin, and 50 mM 2-DG with 4 readings (mix: 3 min; wait: 2 min; measure: 3 min) after each injection. The final values were normalized to cells/well using the CyQUANT Cell Proliferation Assay Kit (Invitrogen, C7026).

### Metabolite quenching and extraction

For tissue specimens, the first step is to deep-freeze in liquid nitrogen. Tissue samples were cryopulverized into fine powder by manual grinding in a mortar and pestle at cold temperature (dry ice). Roughly 2-5 mg of tissue is sufficient for liquid chromatography/mass spectrometry (LC-MS) analysis. For polar metabolite extraction, all extraction solvents must be LC-MS-grade reagents. Roughly 3 mg of tissue from each sample was mixed with 1000 µL extraction solution (600 µL of 60% methanol + ISTD and 400 µL of chloroform). After vortexing continuously for 10 mins at 4°C, the extracted samples were spun down at 4°C, 13.3 krpm, for 15 mins. The metabolite extracts were dried in a vacuum concentrator at 4°C, then resuspended to a final concentration of 30 mg/mL in the aqueous and organic fractions. Lastly, the resuspended metabolite extracts were run on a Thermo Q Exactive Orbitrap Mass Spectrometer for both targeted and untargeted metabolomics.

### Chromatography separation

The metabolite extracts were separated using an iHILIC-(P) Classic column (2.1 µm, 150 mm 3 2.0 mm I.D., The Nest Group) coupled to a Thermo Scientific SII UPLC system. The autosampler and column oven were held at 4 °C and 25 °C, respectively. The iHILIC-(P) Classic column was used with buffer A (0.1% ammonium hydroxide, 20 mM ammonium carbonate) and buffer B (100% acetonitrile) for a representative cross-section of major carbon and nitrogen-handling pathways. The chromatographic gradient was run at a flow rate of 0.150 ml/minute as follows: 0–20 minutes: linear gradient from 80% to 20% B; 20–20.5 minutes: linear gradient from 20% to 80%B; 20.5–28 minutes: hold at 80% B.

### Mass spectrometry and data quality control

The polar metabolites were resolved on a Vanquish U-HPLC system, and detection was performed on a Q Extractive HF-X hybrid quadrupole-orbitrap mass spectrometer (Thermo Fisher Scientific) operated in full-scan mode in both positive and negative ion modes. For metabolite quantification, TraceFinder software (Thermo Fisher Scientific) was used. The lower phase after chloroform extraction was also dried down in a Vacufuge Plus speed-vac at 4 °C, and proteins were resuspended in 0.2 M NaOH and heated at 90 °C for 15 minutes. The protein concentration was determined using the Pierce BCA Protein Assay Kit (Thermo Fisher Scientific, cat. 23227). The metabolite levels were normalized to the total protein amount per sample (mg).

### Data processing and metabolite identification

Peak area integration and metabolite identification were done by using accurate mass and retention time curated with in-house standard library compounds. Data was normalized by tumor weight and/or protein amount and was further normalized by internal standard (D8-phenylalanine) that was spiked during metabolite extraction to account for variations introduced during sample handling, preparation, and injection. The total ion chromatogram (TIC) for each sample was used in addition to the tumor weight for additional normalization. MS data acquisition and targeted feature extraction and quantification were performed using TraceFinder v4.1 (Thermo Fisher Scientific). Untargeted analysis was performed using Compound Discoverer v.3.3, where data-dependent acquisition of fragmentation mass spectra (MS2 data) was used to identify unknowns against the largest, most curated mzCloud spectral library.

### Stable isotope tracing

Cells were grown in triplicate in 20 cm dishes for 24 h in complete medium. The next day, cells were washed twice with PBS, and the medium was removed and replaced with conditional glucose-free RPMI medium supplemented with 10% dialyzed FBS (GIBCO), 1% penicillin, and 4.5 mg/ml ^13^C_6_-D-Glucose (Cat#. CLM-1396-1, Cambridge Isotope) for glucose tracing. For glutamine tracing, cells were washed with PBS, and the medium was removed and replaced with conditional glutamine-free RPMI medium supplemented with 10% dialyzed FBS (GIBCO), 1% penicillin, and 2 mM ^13^C_5_-glutamine (CLM-1822-H-0.1, Cambridge Isotope) and 2 mM ^15^Nγ-glutamine (NLM-557-1, Cambridge Isotope). Specifically, for the untargeted metabolomics experiment with ^13^C_6_-D-Glucose labeling in Fig. 4C, cells were incubated in glucose-free medium containing 10% dialyzed FBS and 4.5 mg/ml ^13^C_6_-D-Glucose for 4-8 hrs. For untargeted metabolomics with ^13^C_5_-glutamine labeling in Fig. 5B-E and ^15^Nγ-glutamine labeling in Fig. 5G, cells were incubated with glucose-free medium with 10% dialyzed FBS containing 2 mM ^13^C_5_-glutamine and 2 mM ^15^Nγ-glutamine for 24 hrs, respectively. Then, metabolites were extracted and detected via LC-MS. Metabolite extraction and metabolomics were performed as described (69). Metabolite peak detection, alignment, identification, and quantification, as well as tracing the incorporation of the 13C label into various metabolites, were processed using Compound Discoverer v.3.3 software (Thermo Fisher Scientific). Peak areas were normalized to protein quantification measured by the Lowry protein assay. For data analysis, statistical significance was assessed using a two-way analysis of variance (ANOVA). The results were considered statistically significant when the p-value was < 0.05. All statistical tests were performed using GraphPad Prism v.10 software.

### Peptides extraction

Cryo-pulverized tissue powders were lysed in 500 µL of lysis buffer (75 mM sodium chloride, 10 mM sodium pyrophosphate, 10 mM sodium fluoride, 10 mM β-glycerophosphate, 10 mM sodium orthovanadate, 50 mM EPPS, pH 8.5, cOmplete Mini EDTA-free protease inhibitor cocktail, 3% SDS, and 5 mM PMSF). Disulfide bonds were reduced by adding dithiothreitol (DTT) to a final concentration of 5 mM and incubating at 56 °C for 30 min. Samples were then alkylated with iodoacetamide (IAA) to a final concentration of 15 mM and incubated in the dark at room temperature for 20 min. Excess IAA was quenched by adding DTT to a final concentration of 5 mM and incubating in the dark at room temperature for 20 min. Proteins were precipitated by adding trichloroacetic acid (TCA; 1 part TCA to 4 parts protein solution) and incubating on ice for 10 min. Precipitated proteins were pelleted by centrifugation (15,000 × g, 10 min, 5 °C) and washed twice with prechilled acetone (−20 °C; 300 µL; 15,000 × g, 10 min, 5 °C). Pellets were resuspended in 500 µL of 1 M urea in 50 mM EPPS (pH 8.5) containing 10 mM calcium chloride, and digested overnight at room temperature with endoproteinase Lys-C (Wako; 1 µg/µL), followed by digestion with sequencing-grade trypsin (Promega; final concentration 1 ng/µL) for 6 h at 37 °C. Digestion was quenched with 1% trifluoroacetic acid (TFA), and peptides were desalted using Sep-Pak C18 solid-phase extraction cartridges (Waters). Peptide concentrations were determined by BCA assay (Thermo Scientific).

### TMT labeling and peptide fragmentation

50 µg of peptides were dried and resuspended in 50 µL of dried and resuspended in 50 µL of 200 mM EPPS (pH 8.5) and 30% acetonitrile (ACN). TMT labeling was performed by adding , 325 µg TMT reagent (Thermo Scientific) in anhydrous ACN, and incubating at RT for 1 h. The reaction was stopped by adding 5% (w/v) hydroxylamine in 200 mM EPPS (pH 8.5) to a final concentration of 0.5% hydroxylamine and incubating at RT for 15 min. Samples were acidified with 1% TFA, combined, desalted over Sep-Pak C18 SPE cartridges, and fractionated by basic pH reversed-phase HPLC (HPRP) (70). All fractions were resuspended in 7.5 uL 5% ACN/5% formic acid and analyzed in 3-hour runs via reversed-phase LC-MS2/MS3 on an Orbitrap Fusion mass spectrometer using the Simultaneous Precursor Selection (SPS) supported MS3 method (71-73). The analysis was performed in an MS1 scan from 500 to 1,200 m/z using the Orbitrap analyzer at a resolution of 6x10^^4^, automatic gain control (AGC) of 5x10^^5^, and a maximum injection time of 100 ms. Fragment ions were subjected to MS2 scans based on abundance, and MS2 and MS3 scans were done within a 5-second cycle. The isolation window was set to 0.5 m/z. Peptides were fragmented using CID at 30% normalized collision energy at the rapid scan rate using an AGC target of 1x10^4^ and a maximum ion injection time of 35 ms using the ion trap. MS3 analysis was performed using synchronous precursor selection (SPS), which simultaneously isolated up to 6 fragment ions.

### Peptide identification and data processing

MS2 spectra were assigned using an in-house-built proteomics analytical platform (74) and a decoy-database-based search strategy to filter for a false-discovery rate (FDR) of less than 1% for peptide and protein assignments (75). We then used linear discriminant analysis (75) for peptide assignment filtering and further calculated the odds of incorrect assignment from a posterior error histogram. Protein assignments were further sorted based on their odds of incorrect assignment, and decoy database matches were used to finalize filtering to an FDR of less than 1%. Peptides that matched more than one protein were assigned to the protein containing the largest number of matched redundant peptide sequences, following the law of parsimony (74). MS3 TMT reporter ion intensities were extracted from the most intense ion within a 0.003 m/z window centered at the predicted m/z value for each reporter ion. Spectra were used for quantification if the average S/N value for all TMT channels was ≥ 40 and the isolation specificity (71) for the precursor ion was ≥ 0.75. Protein intensities were calculated by summing the TMT reporter ions for all peptides assigned to a protein. Intensities were first normalized by the average intensity across all TMT channels relative to the median average across all proteins. In a second normalization step, protein intensities were normalized to the median protein intensity across samples (73).

### Ingenuity Pathway Analysis

To explore overrepresented pathways, the differentially expressed metabolites and proteins (by gene name) were uploaded into Ingenuity Pathways Analysis (IPA) software for core analysis. A p-value of 0.05 was used as the cut-off for determining statistically significant pathway enrichment. The Z-score was used as an indicator to predict activation or inhibition. The reference library was using the Ingenuity Knowledge Base.

### Gene Set Enrichment Analysis

Data were downloaded from cBioPortal for Gene Set Enrichment Analysis (GSEA). Breast tumors were divided into two groups: one with SIRT5 diploidy and one with SIRT5 gain/ amplification. A gene expression matrix was generated for each group, where rows represent genes and columns represent SIRT5 diploidy, gain, and amplification. Genes were ranked based on a metric quantifying their correlation with SIRT5 copy-number alterations using Limma in R (https://doi.org/10.1093/nar/gkv007). Limma was used to perform differential gene enrichment analysis, and genes were ranked based on Sign(log2FC)* -log10(padj) and sorted in descending order. Finally, we used the *GSEApy Python package* to run GSEA with the 2024 Hallmarks Gene Set. Gene sets with a false discovery rate < 0.05 were considered significant.

### TCGA and METABRIC Data Mining

The Tumor Cancer Genome Atlas (TCGA) and the Molecular Taxonomy of Breast Cancer International Consortium (METABRIC) breast cancer data were downloaded from the cBioPortal (https://www.cbioportal.org/study/summary?id=brca_tcga_pan_can_atlas_2018) and (https://www.cbioportal.org/study/summary?id=brca_metabric), respectively. An inner join between mrna_seqv2 and clinical_patient was used to generate SIRT5 mRNA expression for each SAMPLE_ID. Then, the expression was transformed upon log2(SIRT5 expression + 1) transformation. The differences in SIRT5 mRNA expression in breast cancer patients were determined using a t-test comparing BRCA_Basal with other subtypes, as implemented in scipy.stats.ttest_ind(). The calculated CNA frequency for each CNA-PAM50 combination was obtained by inner joining the CNA and the clinical patient data to obtain the SIRT5 CNA and PAM50 subtype for each SAMPLE_ID. A t-test was performed to determine the differences between AMP/GAIN vs. Diploid and AMP/GAIN vs Shallow/Deep Deletion using SciPy Python package.

### cBioportal CPTAC Data Mining

The CPTAC data were downloaded from the cBioPortal (https://www.cbioportal.org/ <u>study/summary?id=brca_cptac_2020</u>). Breast tumors were divided into two groups: SIRT5 amplification/gain and SIRT5 diploid. Any samples with duplicated sample IDs were dropped. The differential expression analysis was performed using Limma in R (https://doi.org/10.1093/nar/gkv007). The upregulated proteins were defined by padj < 0.05 and log2FC > 0.585, and the down-regulated proteins were defined by padj < 0.05 and log2FC < -0.585. The volcano plot was created by Plotly Python package.

### DepMap CRISPR Screening

The DepMap Data Explorer 2.0 was used to investigate the genetic dependencies underlying SIRT5 mRNA expression across 50 breast cancer cell lines via CRISPR screening in the DepMap Portal. The CRISPR data (DepMap Public 25Q2 + Score, Chronos) were analyzed alongside SIRT5 expression data from the Expression Public 25Q2 database. To generate Fig. 7A, the average gene dependency score for each gene effect was computed across the cell lines. Pearson correlation analysis was performed to assess relationships among features across breast cancer cell lines, calculating corresponding p-values and q-values. To visualize the genetic dependency of ATR on SIRT5 expression, the effects of the ATR gene were correlated with SIRT5 mRNA expression in both the breast cancer cohort and across all cancer cell lines. SIRT5 mRNA expression was transformed using the log2 (TPM + 1) method, and Pearson’s correlation was utilized for statistical evaluation.

### Kaplan-Meier Analysis

Relapse-free survival was analyzed by comparing SIRT5 high versus SIRT5 low expression in basal-like breast cancer, stratified by PAM50 subtypes. Breast cancer patients were split by SIRT5 median expression using the GEO database in K-M plotter (76).

### Statistical analysis

Results represent the Mean ± SEM (standard error of the Mean) for the indicated experiments with at least 3 independent biological replicas. The unpaired two-tailed Student’s t-test and the Wilcoxon test were used for comparing two groups, with significance set at a p-value of less than 0.05. GraphPad Prism 10 software was used for statistical analysis. Part of the computational analyses and graph plotting were done in R (version 4.2.0). Statistical analysis of metabolomics data was performed using the MetaboAnalyst platform with parameter settings including no data filtering, log transformation to base 10, and data autoscaling. The Metaboverse application is used to visualize regulatory pattern searches, nearest-neighbor searches, and prioritization (38).

## Supplemental Tables and Figures

**Sup Table 1-2.**
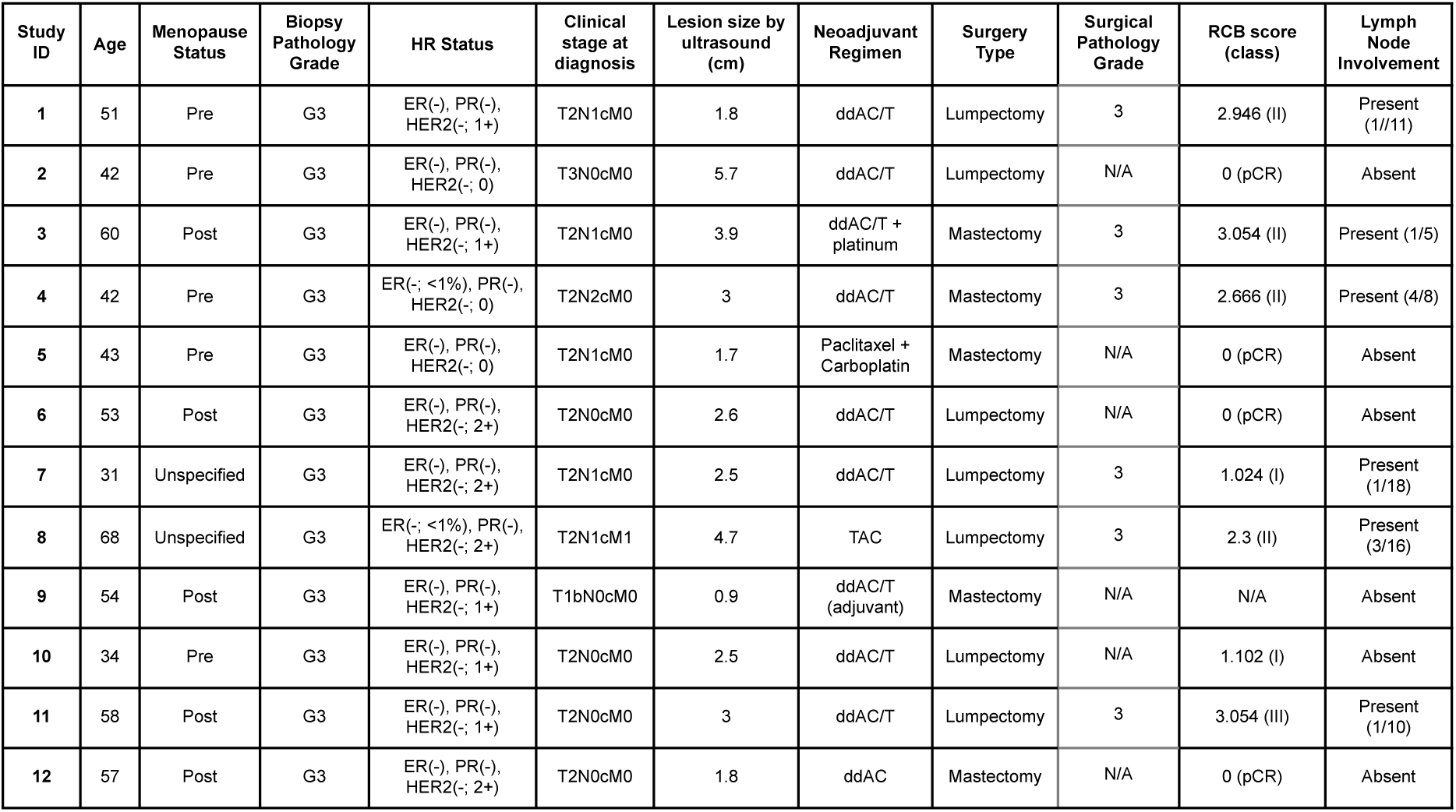
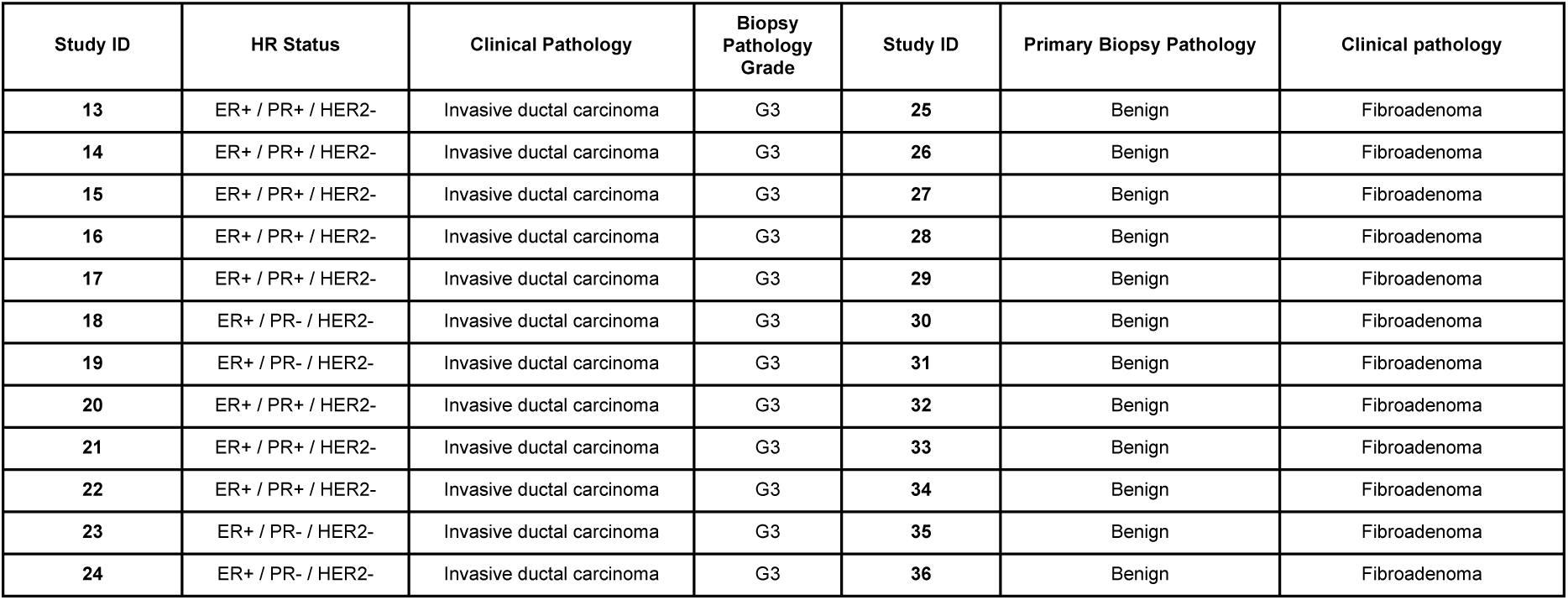
Summary of clinical characteristics of breast tumor biopsies. Clinical data were presented for 12 TNBCs, 12 HR+ IDC, and 12 benign fibroadenomas per the 2018 ASCO/CAP guidelines. HER2 status classification is based on IHC staining, fluorescence in situ hybridization (FISH), and HER2 copy number assessment per cell. TNBC patients who exhibit a non-pCR with residual cancer burden (RCB) are categorized into RCB classes I-III. Abbreviation note: HR, hormonal receptor; ER, estrogen receptor; PR, progesterone receptor; HER2, human epidermal growth factor receptor 2; TNM, tumor-node-metastasis; ddAC/T, dose-dense adriamycin (doxorubicin) and cyclophosphamide followed by taxol (paclitaxel); TAC, paclitaxel, adriamycin, and cyclophosphamide; pCR, pathological complete response; RCB, residual cancer burden; n/a, not applicable.

**Sup Fig. 1.**
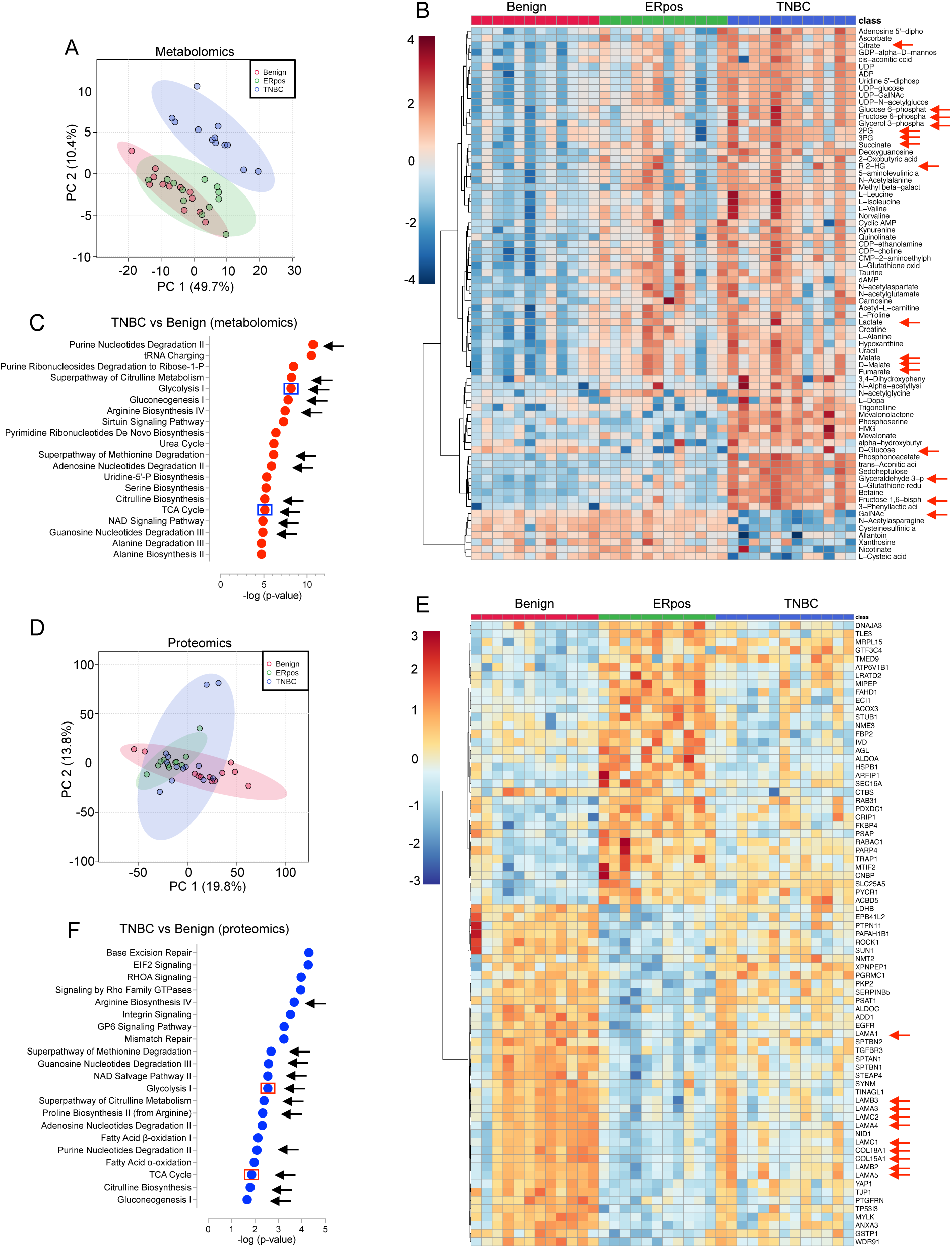
Metabolomic and proteomic profiling reveals metabolic heterogeneity in TNBC. **A.** Principal component analysis (PCA) of metabolites (n=155) from primary TNBC tumors (n=12), HR+ breast tumors (n=12), and benign breast fibroadenomas (n=12) illustrates distinct clustering patterns. **B.** Heatmap clustering of the top significant 75 metabolites across TNBC primary tumors, HR+ breast tumors, and benign fibroadenomas reveals differential expression. Hierarchical clustering is performed using the hclust function in the MetaboAnalyst software (version 6.0); statistical significance is assessed via ANOVA; certain metabolites are highlighted with arrows. **C.** Ingenuity pathway analysis (IPA) shows enriched canonical pathways in TNBCs upon the differentially expressed metabolites between TNBCs and benign lesions; Arrows highlight pathways that are recapitulated in proteomic analysis, and glycolysis and the TCA cycle are particularly highlighted. **D.** PCA of proteins (n=3879) profiled from primary TNBC tumors (n=12), HR+ breast tumors (n=12), and benign breast lesions (n=12) demonstrates distinct protein expression patterns. **E.** Heatmap clustering of the top significant proteins (n=75) across the three groups reveals significant expression differences, determined by ANOVA; certain proteins are highlighted. **F.** IPA shows enriched canonical pathways in TNBCs upon the differentially expressed proteins between TNBCs and benign lesions.

**Sup Fig. 2.**
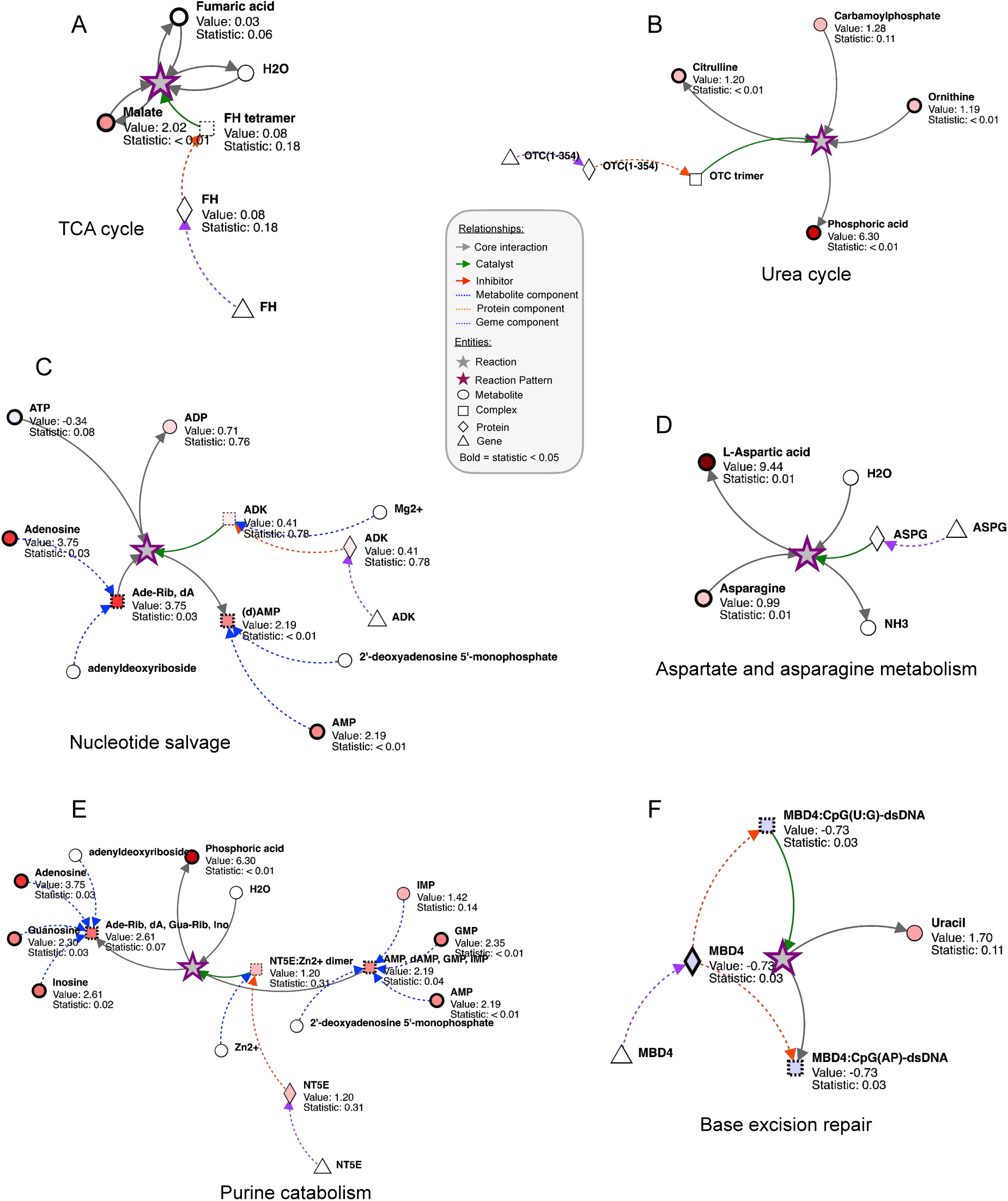
Metaboverse analysis recapitulates the top-enriched metabolic networks identified in IPA. **A-F**. Depictions of six top-ranking, non-redundant collapsed reaction patterns identified by Metaboverse using input from metabolomic and proteomic profiles in non-pCR versus pCR TNBC tumors. Dashed grey edges indicate connections between the distal ends of two to three reactions that have been collapsed. Stars with a solid purple border signify a collapsed reaction. These reactions were identified using the average reaction pattern. **A.** TCA cycle: Fumarate hydratase tetramer hydrolyzes fumarate to malate. **B.** Urea cycle: Ornithine transcarbamylase (OTC) catalyzes the conversion of carbamoyl phosphate and ornithine into citrulline and orthophosphate. **C.** Nucleotide salvage: Adenylate kinase (ADK) catalyzes the conversion of adenosine and ATP to (d)AMP and ADP. **D.** Aspartate and asparagine metabolism: asparaginase (ASPG) hydrolyzes L-Asn to L-Asp. **E.** Purine catabolism: 5’-nucleotidase ecto (NT5E):Zn2+ hydrolyzes AMP, dAMP, GMP, and IMP to produce inosine, guanosine, adenosine, and phosphoric acid. **F.** Base excision repair: MBD4 glycosylase catalyzes the cleavage of uracil. P-values were calculated using a two-tailed, homoscedastic Student’s t-test and were adjusted via the Benjamini-Hochberg correction procedure.

**Sup Fig. 3.**
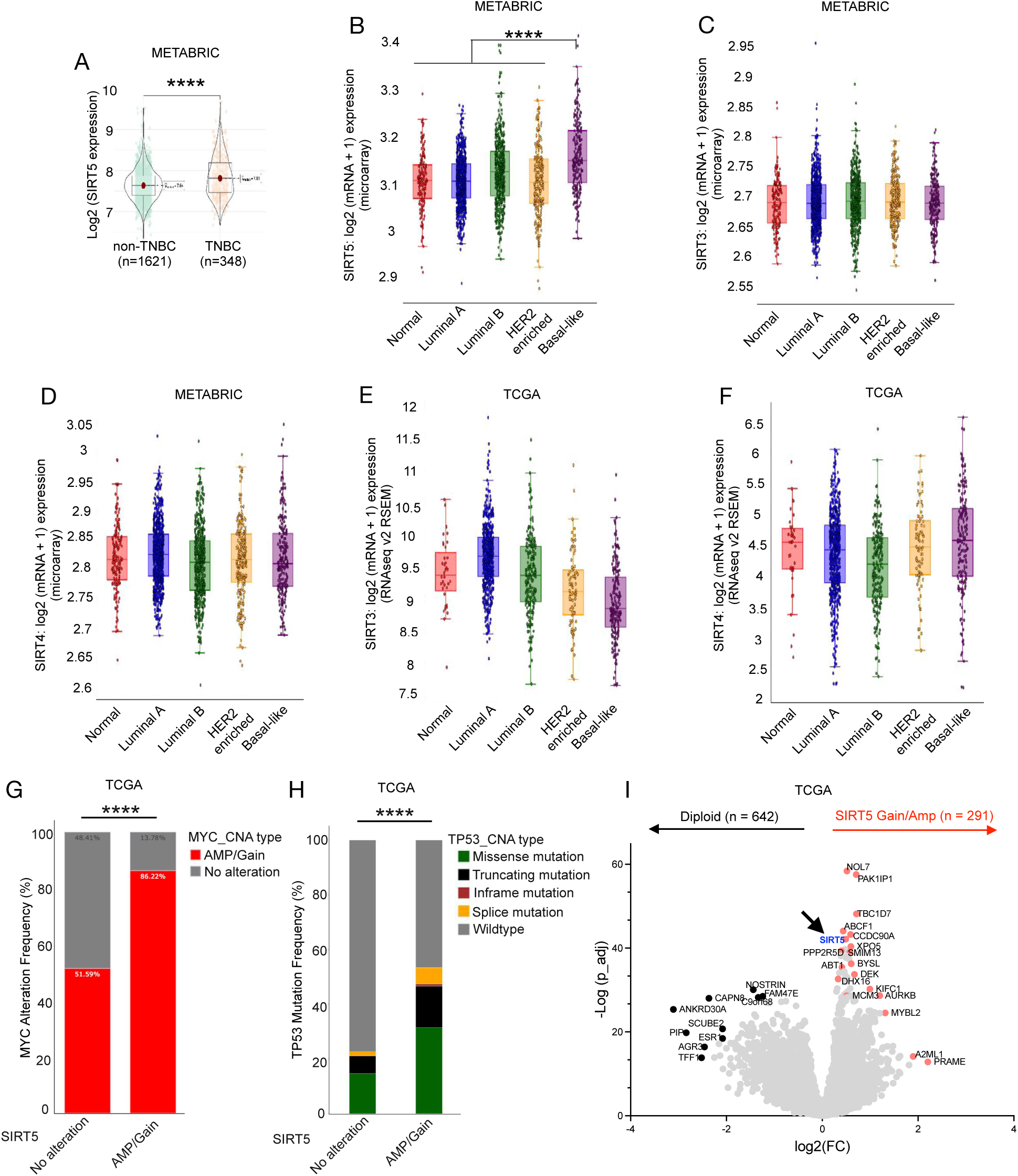
SIR5 is the only mitochondrial sirtuin highly expressed in basal-like breast cancer. **A.** Violin plots illustrate the distribution of SIRT5 mRNA expression levels between TNBC and non-TNBC samples from the Metabric database, revealing a significant increase in expression in TNBC. **B-D**. Box plots depict log2-transformed SIRT3, SIRT4, and SIRT5 mRNA expression levels across breast cancer subtypes classified by the PAM50 gene expression signature from the Metabric database, consistently revealing that SIRT5 is significantly highly expressed in basal-like breast cancer, while SIRT3 and SIRT4 are not differentially expressed in different subtypes of breast cancer. **E-F**. Box plots depict log2-transformed SIRT3 and SIRT4 mRNA expression levels across breast cancer subtypes classified by the PAM50 gene expression signature from the TCGA database, consistently revealing that SIRT3 and SIRT4 are not differentially expressed in different subtypes of breast cancer. **G-H**. Bar graphs show that SIRT5 amplification/gain is significantly associated with MYC amplification and TP53 mutation, particularly missense and truncating mutations. **I.** The volcano plot illustrates the differentially expressed genes in breast tumors with SIRT5 amplification/gain (n = 291) compared to diploid tumors (n = 642) from the TCGA database. The arrow highlights SIRT5 expression in the plot. **** p-value < 0.0001.

**Sup Fig. 4.**
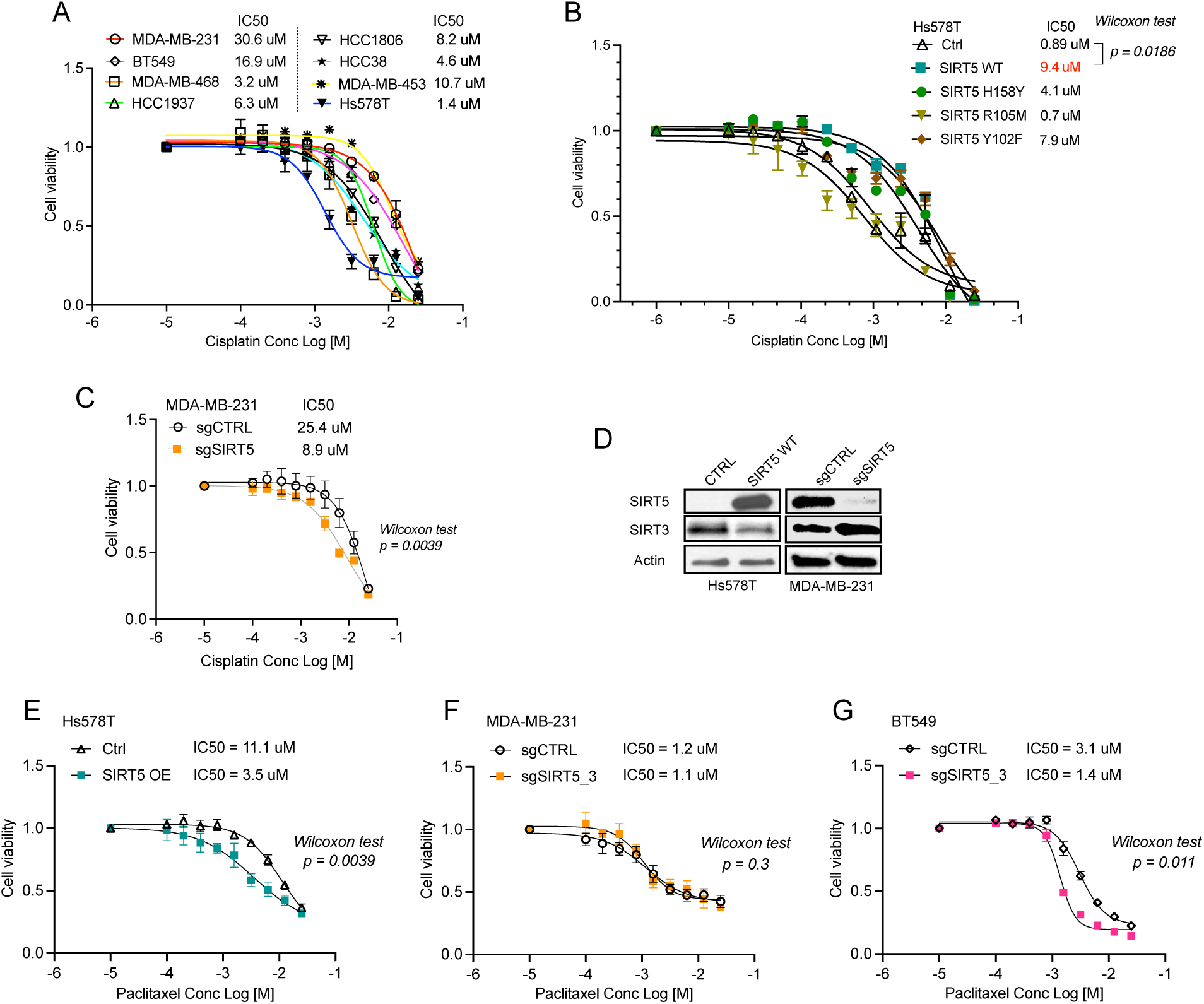
SIRT5 enhances chemoresistance to cisplatin. **A.** The dose-response curves illustrate the sensitivity of several TNBC cell lines to cisplatin, alongside the corresponding IC50 values. **B.** The dose-response curves demonstrate the sensitivity of Hs578T cells expressing control vector, SIRT5 wildtype, and SIRT5 mutants (H158Y, R105M, and Y102F) to cisplatin, accompanied by their respective IC50 values. **C.** The dose-response curves illustrate the response of MDA-MB-231 cells to cisplatin in the presence or absence of SIRT5 expression, along with their respective IC50 values. **D.** Western blotting shows SIRT5 and actin levels in Hs578T and MDA-MB-231 cell lines. **E-G**. The dose-response curves illustrate the response of Hs578T, MDA-MB-231, and BT549 cells to paclitaxel, along with their corresponding IC50 values.

**Sup Fig. 5.**
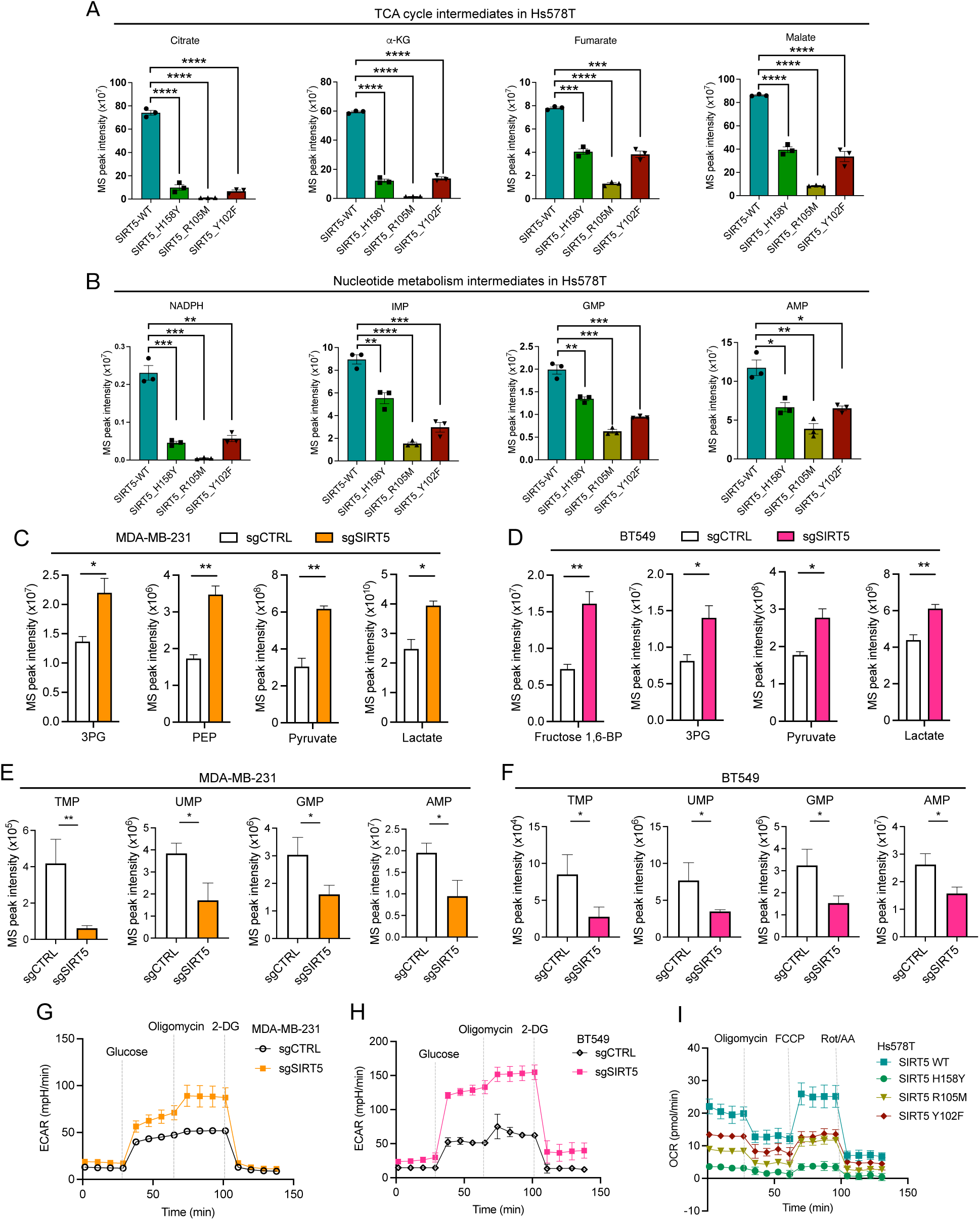
SIRT5 catalytic mutations result in reduced TCA activity and nucleotide biosynthesis. **A-B**. Bar graphs illustrate the MS peak intensity of citrate, α-KG, fumarate, and malate in the TCA cycle, as well as the metabolite abundance of NADPH, IMP, GMP, and AMP in nucleotide metabolism. **C-D**. Bar graphs depict the MS peak intensity of glycolytic intermediates in MDA-MB-231 and BT549 cells under SIRT5 knockout (sgSIRT5) conditions compared to control (sgCTRL) conditions. **E-F**. Bar graphs showing the MS peak intensity of nucleotide metabolites (TMP, UMP, GMP, and AMP) in MDA-MB-231 and BT549 cells under scrambled sgCTRL compared to SIRT5 knockout conditions, indicating a significant decrease in nucleotide levels associated with SIRT5 loss. **G-H**. Glycolysis stress test measures ECAR in MDA-MB-231 and BT549 cells in the presence or absence of SIRT5 expression. **I.** Mitochondrial stress test measures OCR in Hs578T cells expressing wildtype SIRT5 (SIRT5 WT) versus three SIRT5 mutants (H158Y, R105M, and Y102F). ns indicates non-significance; statistical significance is indicated with * p-value < 0.05; ** p-value < 0.01; *** p-value < 0.001; **** p-value < 0.0001.

**Sup Fig. 6.**
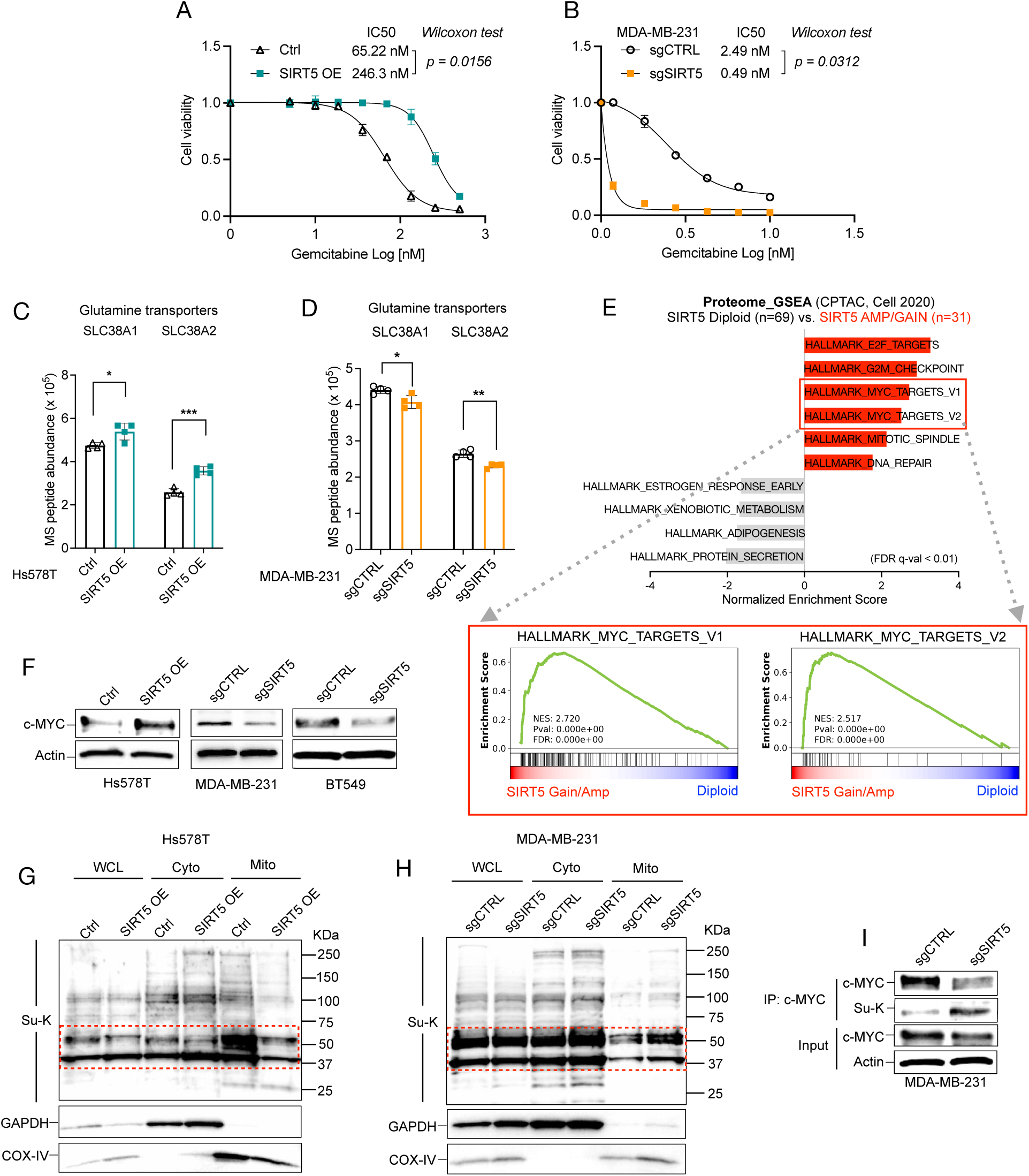
SIRT5 facilitates glutamine catabolism to sustain TCA cycle anapleurosis through the activation of cMYC. **A-B**. The dose-response curves illustrate the sensitivity of Hs578T and MDA-MB-231 cells to gemcitabine in the presence or absence of SIRT5 expression, along with their corresponding IC50 values. **C-D.** Bar graphs show the peptide abundance detected by MS for glutamine transporters, including SLC38A1 and SLC38A2, in Hs578T and MDA-MB-231 cells with or without SIRT5 expression. **E.** GSEA on the proteome of breast tumors (n = 100) sourced from the cBioPortal CPTAC database identifies hallmark pathways that are significantly upregulated and downregulated in tumors with SIRT5 gain/amplification (n = 31) compared to diploid tumors (n = 69). **F.** Western blotting shows cMYC and actin protein levels in Hs578T, MDA-MB-231, and BT549 cells with or without SIRT5 expression. **G-H**. Western blotting measures the levels of succinylated lysine (Su-K) in whole cell lysate (WCL), cytosolic proteins (Cyto), and mitochondrial proteins (Mito) in Hs578T (G) and MDA-MB-231 (H) cells. Each panel includes quantification of Su-K, with COX-IV and GAPDH used as loading controls. The molecular weights of relevant proteins are indicated in KiloDaltons (kDa). **I.** IP Western blotting shows MYC, Su-K, and actin protein levels in MDA-MB-231 cells in the presence or absence of SIRT5 expression. Statistical significance is indicated with * p-value < 0.05; ** p-value < 0.01; *** p-value < 0.001;

**Sup Fig. 7.**
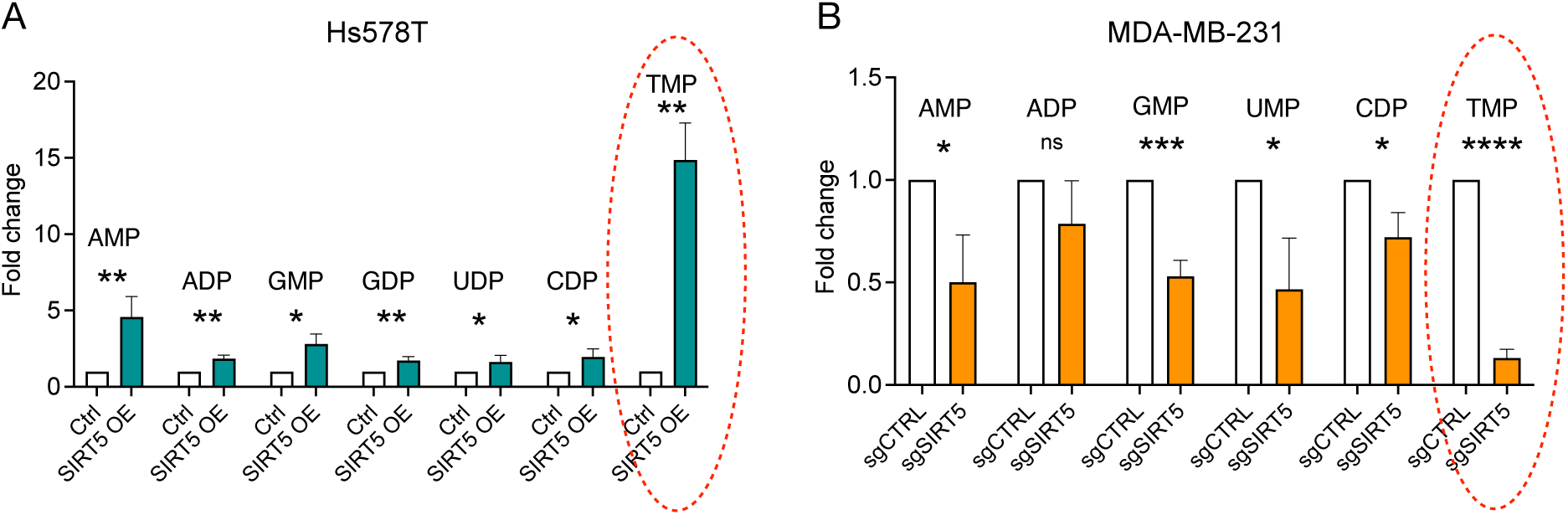
SIRT5 expression selectively enhanced pyrimidine biosynthesis. **A-B**. Bar graphs display the fold changes in nucleotide levels between Hs578T SIRT5 OE and control cells, as well as between MDA-MB-231 sgSIRT5 and sgCTRL cells. ns indicates non-significance; statistical significance is indicated with * p-value < 0.05; ** p-value < 0.01; *** p-value < 0.001; **** p-value < 0.0001.

